# Evidence for a unitary structure of spatial cognition beyond general intelligence

**DOI:** 10.1101/693275

**Authors:** Margherita Malanchini, Kaili Rimfeld, Nicholas G. Shakeshaft, Andrew McMillan, Kerry L. Schofield, Maja Rodic, Valerio Rossi, Yulia Kovas, Philip S. Dale, Elliot M. Tucker-Drob, Robert Plomin

## Abstract

Performance in everyday spatial orientation tasks (e.g. map reading and navigation) has been considered functionally separate from performance on more abstract object-based spatial abilities (e.g. mental rotation and visualization). However, evidence remains scarce and unsystematic. With a novel gamified battery, we assessed six tests of spatial orientation in a virtual environment and examined their association with ten object-based spatial tests, as well as their links to general cognitive ability (*g*). We further estimated the role of genetic and environmental factors in underlying variation and covariation in these spatial tests. Participants (*N* = 2,660) were part of the Twins Early Development Study, aged 19 to 22. The 6 tests of spatial orientation clustered into a single ‘*Navigation’* factor that was 64% heritable. Examining the structure of spatial ability across all 16 tests, three factors emerged: *Navigation, Object Manipulation* and *Visualization.* These, in turn, loaded strongly onto a general factor of *Spatial Ability*, which was highly heritable (84%). A large portion (45%) of this high heritability was independent of *g*. The results from this most comprehensive investigation of spatial abilities to date point towards the existence of a common genetic network that supports all spatial abilities.

---

Spatial skills are fundamental for everyday life as they make it possible for us to understand and operate on the physical world around us. Studies in primates and other animals have highlighted the importance of spatial ability for evolution and survival. Food-hoarding birds rely on spatial memory to retrieve their caches, which is crucial to their subsistence, and climate harshness has been found to positively drive the evolution of spatial memory skills in black-capped chickadee, another bird species.^1^ Spatial skills are also important in modern technologically-oriented societies^2^ as individual differences in spatial skills are associated with positive developmental, educational and life outcomes. Spatial ability reliably predicts scholastic and professional success and career choices, particularly in Science, Technology, Engineering and Mathematics (STEM) and related fields, even after controlling for general cognitive ability.^3–5^ In spite of the increasingly fundamental role that spatial ability has for individuals and contemporary societies,^6^ numerous questions remain regarding the nature of spatial ability as well as its origins and structure.^7^

What constitutes good spatial skills? Since its earliest conceptualization,^8^ spatial ability has been considered a multifaceted construct comprising several related, yet separable, skills.^9^ One of the most widely adopted definitions of spatial ability describes it as the ability to generate, retain, retrieve and transform well-structured visual images.^10^ Contrary to this very broad characterization of spatial ability, however, extant research has largely focused on measuring only specific aspects of object-based spatial ability. Among the most widely studied spatial skills are individuals’ abilities to mentally rotate shapes,^11^ to visualize objects from different perspectives, and to find figures embedded within other shapes.^12^ A much smaller body of research has considered larger-scale, practical everyday spatial orientation abilities, such as navigation, map reading and way-finding.

Until recent years, studies of spatial orientation skills had been hindered by the difficulty in measuring navigation and way-finding abilities in real-life settings utilizing rigorous approaches that are standardized across participants. In addition, assessing navigation in the real environment can be highly costly and time consuming and thus unlikely to be scalable to large samples nation-wide or world-wide. Technological advances in the field of virtual reality (VR) provide a novel powerful tool to study individual differences in spatial orientation skills in realistic settings that can be fully controlled and standardized across participants.^13,14^ Studies assessing the validity of measuring navigation skills using VR have observed strong correlations (∼.60) with performance in real world navigation skills.^13,15^ The reliability of assessing spatial abilities in VR is likely to continue increasing as accelerating technological developments provide progressively immersive and realistic tools.

Likely due, at least in part, to such difficulties in assessing multiple spatial orientation skills reliably in large, representative samples, few studies have examined the structure of spatial orientation ability and its association with other spatial skills. More broadly, evidence concerning the nature and factor structure of spatial ability remains mixed, with most studies focusing on differentiating between relatively few measures rather than examining the communalities across a broad range of spatial skills.^7,10,16,17^ In our previous work,^18^ we have shown that a general factor of spatial ability captures a substantial proportion of variance across numerous tests of spatial skills, and that communalities across tests are largely explained by shared genetic variance.^18^ However, one major limitation characterized our previous study: Although we considered ten object-based spatial abilities, including tests of rotation, visualization and scanning abilities, we did not include measures of spatial orientation, such as navigation, map reading and way-finding.

The omission of spatial orientation measures has special theoretical relevance because evolutionary and cognitive theories have pointed to a distinction between the ability to mentally manipulate objects on a small scale (object-based spatial skills) and the ability to orient in large-scale environments (spatial orientation ability).^19–21^ This proposition is partly supported by psychological studies suggesting that the two abilities are influenced by separate cognitive processes and brain structures. For example, in a study of the association between performance in object-based psychometric spatial tests and large-scale spatial learning, partial support was found for a differentiation between these skills. Individual differences in measures of spatial learning (measuring skills such as placing landmarks on a map, intra-route distance estimates and route reversal) were unrelated to variation in object-based spatial tests. However, the ability to learn maze and maze reversal, was found to be related to both object-based tests and spatial learning.^22^ Other studies in the field of cognitive psychology have found evidence for a partial dissociation between object-based tests and large-scale spatial orientation skills.^23–25^

Neuroimaging studies have also provided preliminary converging evidence for the distinction between object-based abilities and spatial orientation skills, suggesting that the two are supported by separate brain networks. Object-based spatial skills, and particularly mental rotation ability, were found to be primarily associated with activation of the parietal lobes.^26^ Conversely, variation in learning and remembering the layout of large-scale spaces has been found to be related to processing in the hippocampus and the medial temporal lobes.^27^

Other theoretical accounts and studies, however, have suggested that object-based and spatial orientation skills might be closely related. For example, theories concerning the evolution of sex differences have argued that individual variation in object-based spatial skills, such as mental rotation, are the product of different selection pressures for large-scale spatial orientation abilities between males and females over evolutionary history,^28,29^ therefore suggesting that the two largely reflect common skills. Empirical evidence also supports the idea of a largely unitary set of abilities. A study of the association between object-based spatial abilities, measured with a limited battery of three psychometric tests, and large-scale spatial orientation skills, measured both in realistic settings and a virtual environment, found a substantial correlation between the two.^15^

The proposition of a unitary set of cognitive processes underlying object-based and spatial orientation skills is consistent with the idea that these are aspects of a more general set of cognitive abilities. It is plausible that at the heart of individual differences in all spatial skills is general cognitive ability, or general intelligence (*g*). *G* is a psychometric construct that emerged at the beginning of twentieth century from observations that almost all cognitive tests correlate moderately and positively.^30^ Individuals performing highly on one cognitive test are also likely to show good performance on other tests of cognitive abilities, and *g* indexes this covariance observed between cognitive measures. Therefore, *g* is thought to represent individual differences in the domain-general abilities to plan, learn, think abstractly, and solve problems that are necessary for successfully completing cognitive tests.^31^

In our previous work on the factor structure of object-based spatial tests, we have shown that individual differences in spatial abilities cluster into a unitary factor, at both the observed and genetic levels, even after accounting for *g*.^18^ Along the same lines, another study found that the association between object-based and spatial orientation abilities was largely independent of verbal ability.^15^ These studies suggest that the coherence of spatial abilities is not simply due to their being part of *g*, but rather inherent in the spatial domain itself. However, neuropsychological evidence contradicts this view. Case studies of patients with neuropsychological impairments suggest that damage to navigation-related structures in humans typically leads to broad memory deficits that are not limited to the spatial domain.^10^

Extant literature is therefore characterized by contrasting theories and evidence with respect to the factor structure and associations between object-based spatial abilities, assessed mostly through psychometric tests, and large-scale spatial orientation skills, assessed both in real settings and VR. The lack of a cohesive account is likely due to a paucity of studies that have investigated the association between object-based and large-scale spatial orientation skills with a sufficiently diverse battery of tests. In addition, to our knowledge, no study to date has investigated their links within a genetically informative framework, testing the hypothesis that a common genetic network, independent of *g*, supports performance in all spatial skills.

The current study addresses these limitations by investigating the structure of spatial ability using two comprehensive online batteries of object-based and spatial orientation skills, administered to a large genetically-informative sample of twins aged 19 to 22. Importantly, we assessed spatial orientation abilities with an innovative gamified battery of six tests measuring navigation, map reading, wayfinding and large-scale scanning and perspective-taking skills set in a virtual environment. The current work has three main aims: First, we examined, for the first time, the factor structure and origins of spatial orientation skills. Second, we investigated the structure and genetic and environmental origins of spatial ability across sixteen tests of object-based and spatial orientation skills. Third, we explored the role that *g* has in unifying individual differences in performance across tests of spatial abilities.

Addressing outstanding questions on the factor structure of spatial ability applying a genetically informative design provides new evidence about its genetic and environmental underpinnings. This new knowledge provides a critical foundation for future advances in the study of individual differences in spatial cognition, impacting several scientific fields, from cognitive psychology to neuroscience, anthropology and evolutionary biology.

## RESULTS

### Individual differences in spatial orientation can be measured reliably in a virtual environment and are moderately heritable

We first assessed whether our newly developed gamified battery set in a virtual environment could effectively capture individual differences in spatial orientation skills in our large sample of twins. Beyond showing good test-retest reliability (average *r* = .74, ranging from .60 to .89, see Methods for information on the reliability of each tests), the six tests –scanning, perspective taking, navigation based on landmarks, navigation following directions, route memorizing and map reading– showed normal distributions, with acceptable values for skewness and kurtosis (Figure 1; Supplementary Table S1). Therefore, our gamified battery was able to discriminate and reliably capture variation in spatial orientation abilities.

**Figure 1.**
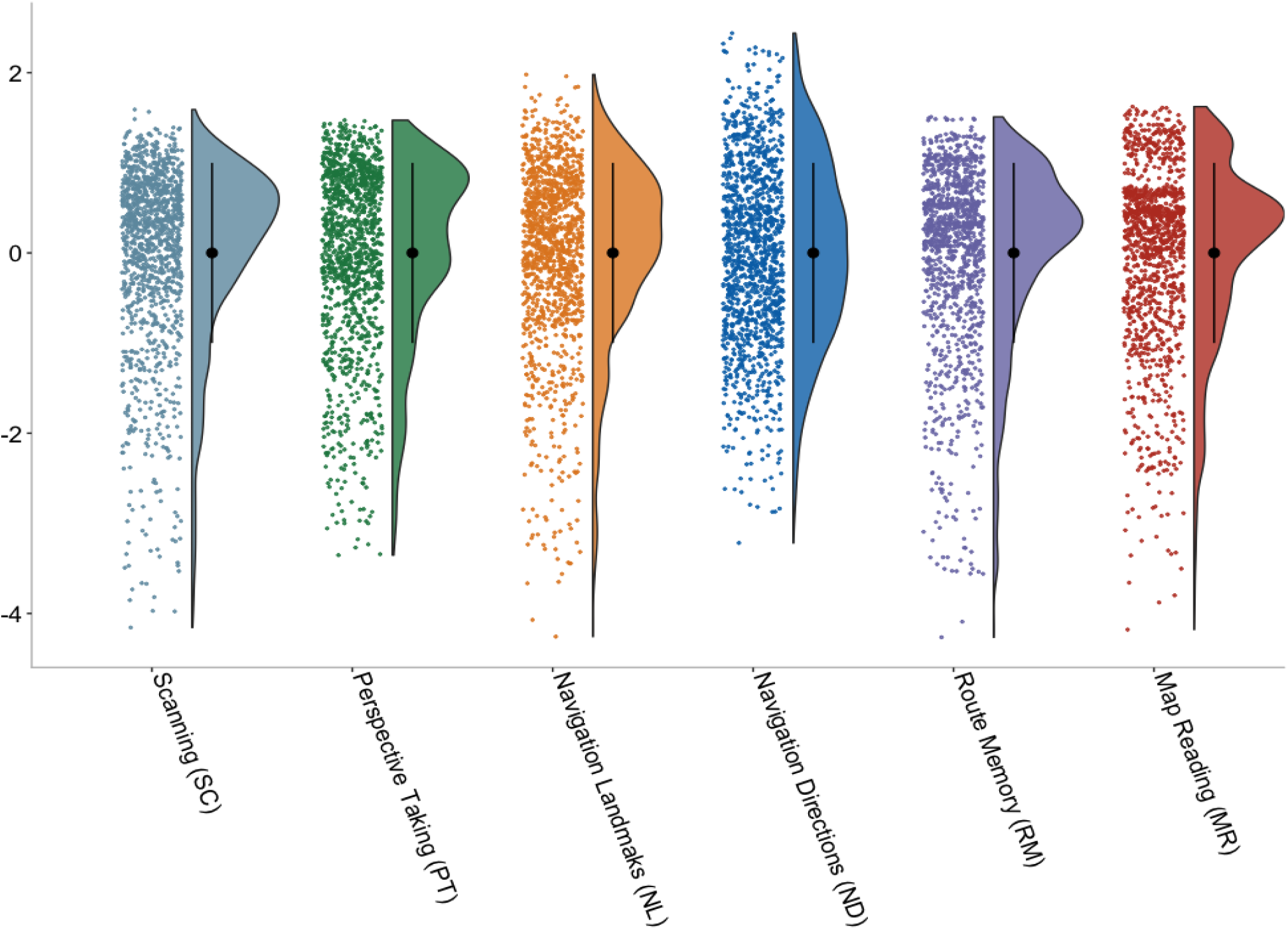
Individual differences and distributions for the six tests included in our novel gamified battery of spatial orientation set in a virtual environment. All variables were residualized for age and sex, and standardized in one randomly-selected half of the sample (only one twin within each pair was randomly selected for descriptive and phenotypic analyses in order to account for the non-independence of observations); full descriptive statistics for both randomly selected halves of the sample are presented in Supplementary Table S1.

We adopted the twin method (Methods) to calculate heritability estimates for the six measures of spatial orientation; these are presented in presented in Figure 2. Heritability estimates, the extent to which variation in a trait is accounted for by genetic differences,^32^ ranged from modest to strong (14-57%). The remaining variance in all tests was accounted for by non-shared environmental factors, environmental factors that do not contribute to similarities between siblings,^32^ with the only exception being the test of orientation ability using landmarks, which showed a significant proportion of shared environmental variance (15%). These substantial non-shared environmental estimates might in part reflect measurement error.

**Figure 2.**
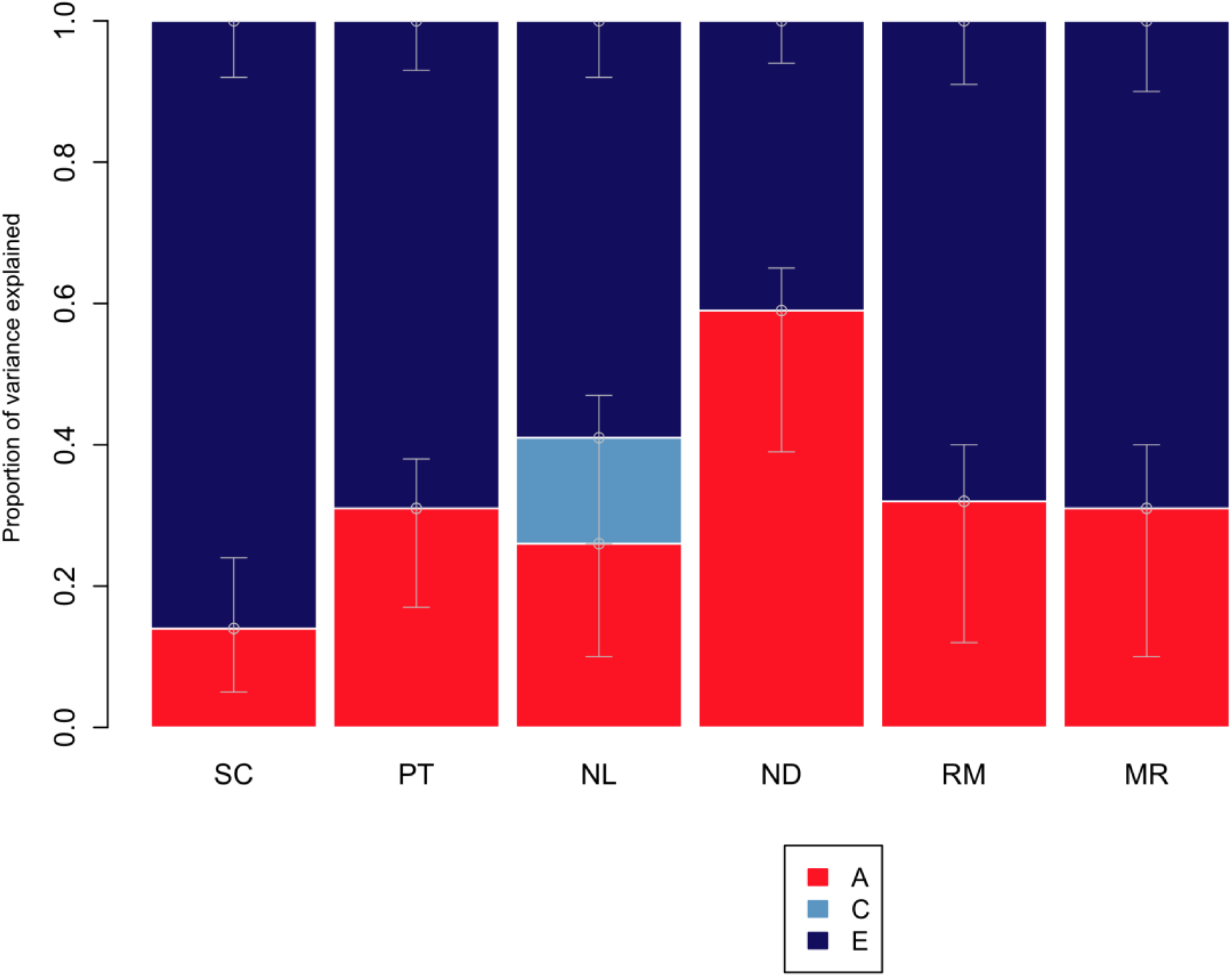
Genetic and environmental estimates for navigation tests: univariate model-fitting results. A: additive genetic; C: shared environmental; E: non-shared environmental components of variance. SC: scanning test; PT: perspective taking; NL: navigation according to landmarks; ND: navigation according to directions (cardinal points); RM: route memory; MR: map reading.

Because sex differences are often found for spatial abilities (though not always in the same direction),^33,34^ we conducted univariate full sex-limitation model (Methods) to examine whether these estimates of heritability differed between males and females. We found no evidence for qualitative genetic sex differences, meaning that the same genetic and environmental factors seemed to influence individual differences in spatial orientation abilities for males and females. No significant quantitative sex differences were found (Supplementary Table S2), that is, differences in the magnitude of genetic and environmental influences. For example, for an overall composite measure of navigation ability, heritability was 52% (95% CI: 0.31; 0.70) for males and 54% for females (95% CI: 0.29; 0.62). These estimates have overlapping confidence intervals, indicating that they are not statistically different from one another. Even with a sample of over 800 complete twin pairs who took part in the spatial orientation battery, the sample size was not sufficient for the sex-limitation model to reliably detect quantitative and qualitative sex differences, if they in fact exist. This is evident from the wide confidence intervals around estimates when calculated for males and females separately. For these reasons, and to increase power, the full sample was used in subsequent analyses, combining males and females, and same- and opposite-sex twin pairs.

### A single ‘navigation’ factor captured the variance common across all tests of spatial orientation

We applied factor analysis (Methods) to examine the covariance structure across the six tests in the spatial orientation battery. The results showed that the six tests correlated substantially and clustered into one common factor, which we named ‘*Navigation*’, as it indexed abilities that are generally described in the literature as spatial navigation skills (see Supplementary Table S3–factor structure, and Figure 4 –intercorrelations between tests). This unifactorial model provided a good fit for the data (χ^2^ = 269.937 (148), *p* = 0.0000; CFI = 0.968; TLI = 0.971; RMSEA = 0.030; SRMR = 0.049).

**Figure 3.**
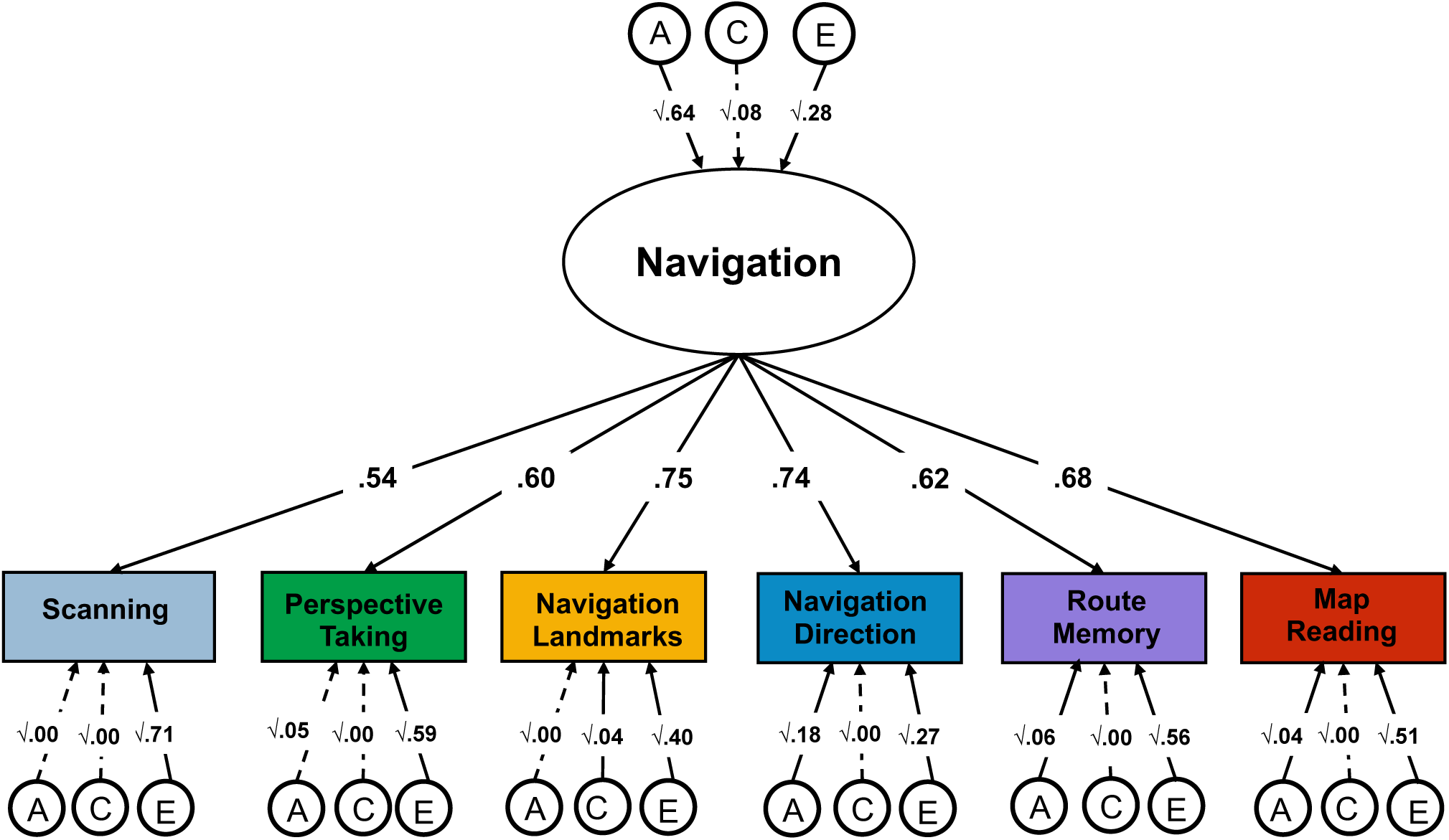
Factor structure and genetic and environmental variance common across the six tests of spatial orientation. We applied the common pathway model to parse the genetic (A), shared environmental (C) and nonshared environmental (E) variance that is shared across all the tests (represented by the A, C and E paths leaving from the common *Navigation* factor) from the genetic and environmental variance that is specific to each test (indexed by the individual A, C, and E latent factors leaving from each rectangle). Each individual test loaded substantially onto a common factor, which we named *Navigation factor* (loadings ranging from λ = .54 for scanning ability to λ = .75 for navigation based on landmarks). All A, C and E paths are standardized and squared.

**Figure 4.**
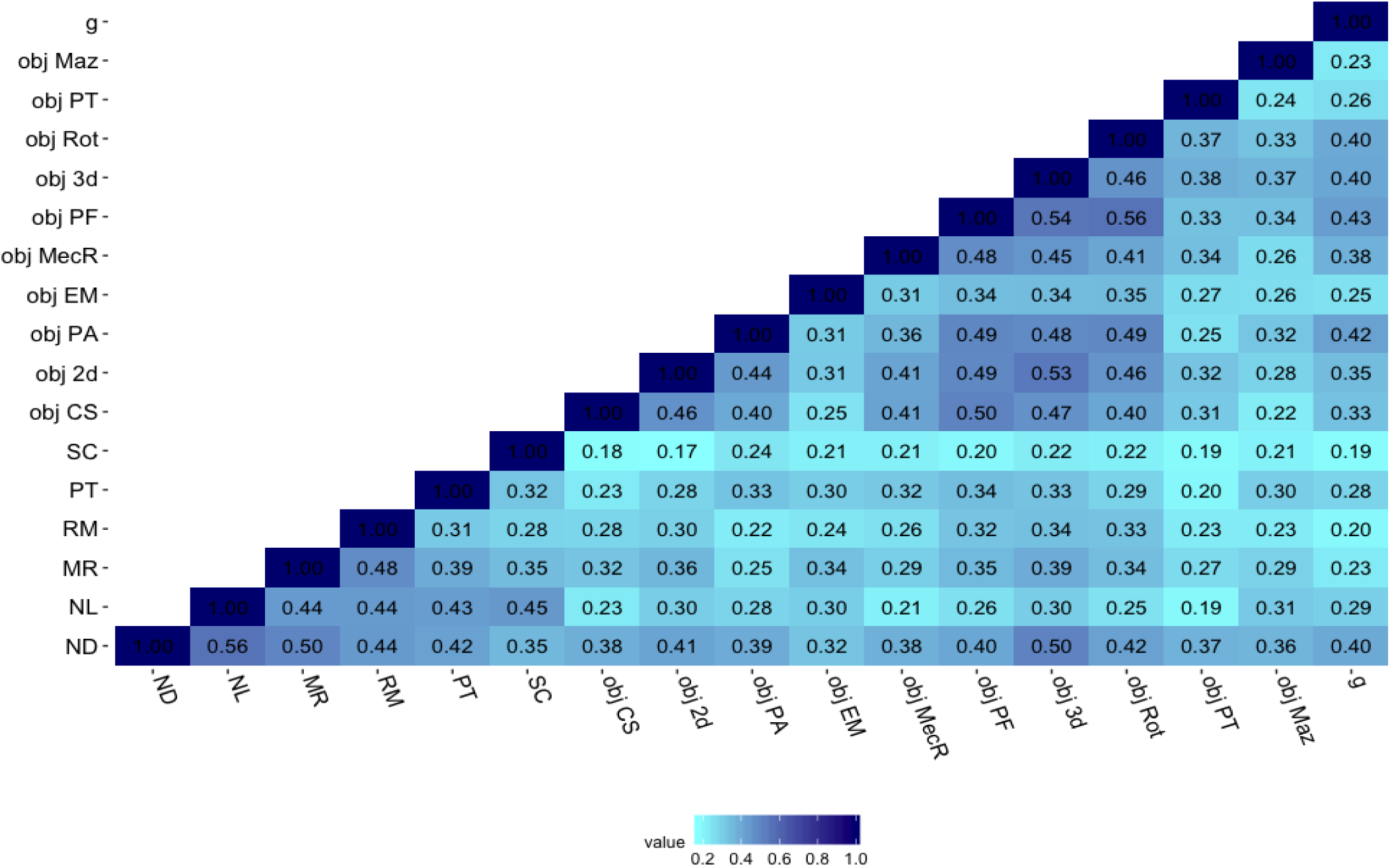
Correlations between the 16 tests of spatial ability and *g*. Starting from the bottom left of the matrix, the first six tests are part of the spatial orientation battery. ND = navigation according to directions, NL = navigation according to landmarks, MR = map reading, RM = route memory, PT = perspective taking, SC = scanning. The following 10 tests were part of the other battery assessing object-based spatial skills: obj CS = cross-section, obj 2d = 2d drawing, obj PA = pattern assembly, obj EM =Elithorn Maze, obj MecR = Mechanical Reasoning, obj PF = paper folding, obj 3d = 3d drawing, obj Rot = mental rotation, obj PT = perspective taking, obj Maz = mazes, *g* = general cognitive ability. All correlations were significant at *p* < .001; variables were residualized for age and sex and standardized prior to analyses.

We used the Common Pathway model (Methods) to examine the extent to which genetic (A), shared environmental (C) and nonshared environmental (E) effects were common or specific across the six tests (Figure 3). We found that the heritability of the common navigation factor was 64% (CIs = 0.41; 0.91); shared environmental and non-shared environmental factors accounted for smaller proportions of variance, 8% (CIs -.00; .43) and 28% (CIs 0.21-0.36), respectively. The largest part of the genetic variance in navigation ability was shared across all tests; between 66% and 100% of the heritability of each test was captured by the common factor of navigation. Consequently, test-specific genetic effects were found to account for between 0% and 34% of the genetic variance in each test of spatial orientation (Supplementary Table S4).

Environmental factors were largely specific to each test, as indicated by the considerable size of the specific E paths (bottom of Figure 3), between 64% and 90% of the nonshared environmental variance was found to be specific to each test. The common navigation factor only captured between 10% and 36% of nonshared environmental variance in each test of spatial orientation (Table S4).

### Substantial associations between measures of spatial orientation and object-based spatial tests

We investigated the structure of spatial ability across a greater diversity of spatial tests. To this end, we extended our analyses beyond the six tests of spatial orientation to incorporate 10 additional tests of object-based spatial skills^18^. This additional battery of spatial tasks included measures that very closely align with traditional psychometric tests of spatial ability, including mental rotation, visualization, 2D and 3D drawing ability, and mechanical reasoning. Figure 4 presents phenotypic correlations between the sixteen spatial tests included in the two batteries (spatial orientation and object-based) and their correlations with *g*.

Correlations between spatial tests were positive and moderate (.17-.56), with stronger links observed between certain tests within each battery. For example, the four tests assessing navigation and map reading skills in the spatial orientation battery clustered more strongly together (*r* ranging from .44 to .56). The same was observed for measures of 2D and 3D drawing, pattern assembly, paper folding, and mental rotation in the object-based battery (*r* ranging from .34 to .54).

We conducted a series of confirmatory factor analyses to formally evaluate the covariance structure between the 16 spatial tests. We tested different theoretical models about the structure of spatial skills, starting from the simplest model and progressing to increasingly complex representations of the structure of spatial skills. The first model we tested was a one-factor model (Figure 5a), positing that variation in spatial orientation and object-based skills could be largely considered a unitary ability. Although all tests loaded substantially onto a single factor (Figure 5a), model fit indices (χ^2^ = 692.730 (104), *p* < .001, CFI = 0.890, TLI = 0.873, RMSEA = 0.061, SMRS = 0.059) suggested that this structure did not provide a good fit for the data.

**Figure 5.**
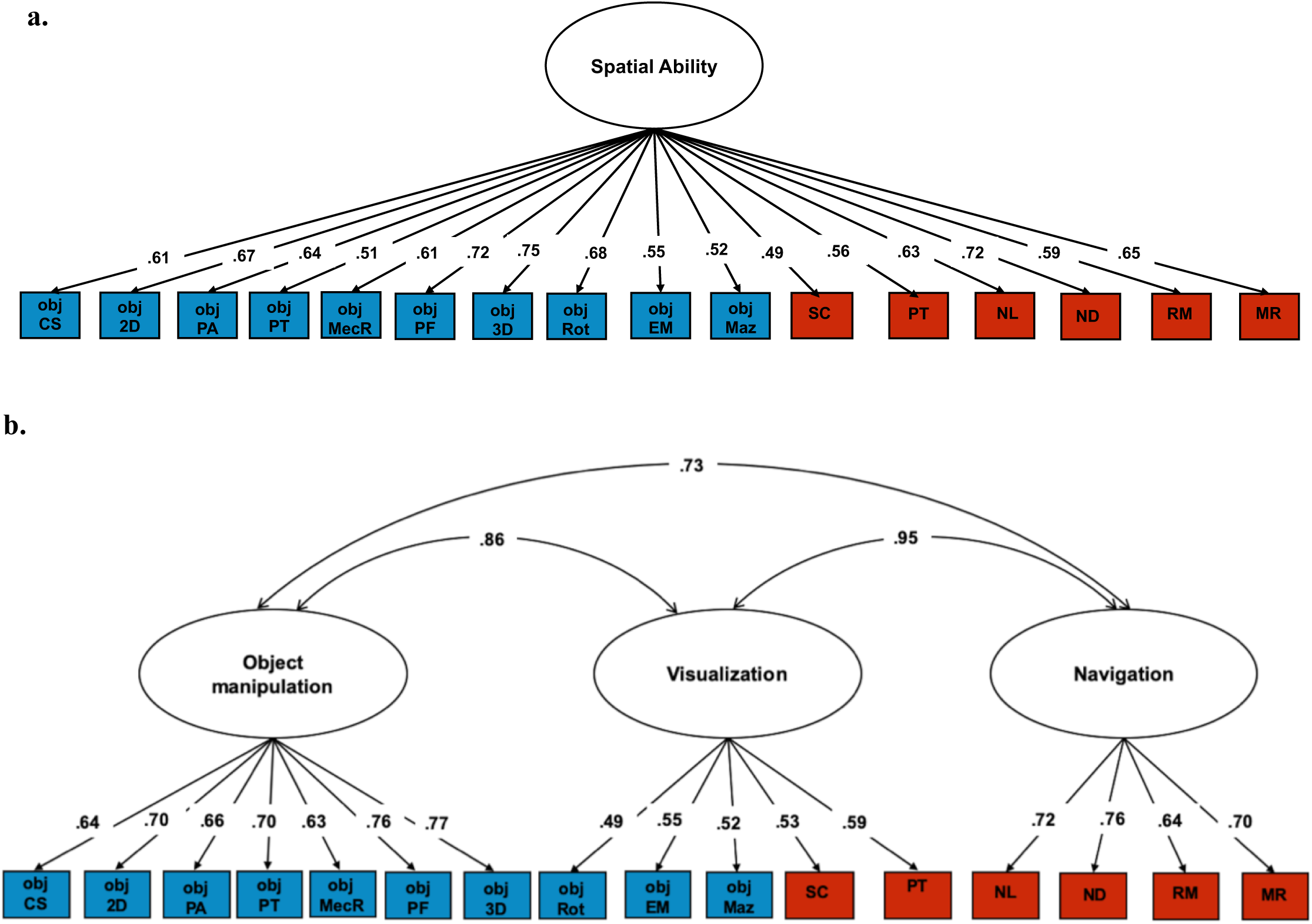
Factor structure across all 16 tests. **(a**) Unifactorial model of spatial ability; **(b)** Three-factor model of spatial ability. Obj CS = cross-section, obj 2D = 2D drawing, obj PA = pattern assembly, obj SR = shapes rotation, obj MecR = mechanical reasoning, obj PF =paper folding, obj 3D = 3D drawing, obj PT = perspective taking, obj EM = Elithorn Maze, obj Maz = Mazes, SC = scanning, PT = perspective taking, NL = navigation according to landmarks, ND = navigation according to directions, RM = route memory, MR = map reading.

Secondly, we tested whether including two factors of spatial ability (one for each battery, Figure S1) would provide a more accurate description of the structure of spatial skills. This model provided a good fit (χ^2^ = 316.000 (103), *p* < .001, CFI = 0.958, TLI = 0.951, RMSEA = 0.037, SMRS = 0.040). However, it also presented one major limitation: due to the substantial difference in test administration and properties of the two batteries, we could not exclude the possibility that the two separate factors emerging from this analysis were a product of differences between the two batteries, rather than underlying a real set of separate, although substantially correlated, abilities. In addition, the two batteries included some cases of parallel measures, so that specific skills were tested in both batteries using different methods (e.g. scanning and perspective taking).

In order to overcome this limitation, we tested another two-factor model, but this time we constructed the two factors based on theoretically-driven differences between the constructs. The first factor included all those tests that are described in the literature as tapping spatial orientation abilities (navigation, way-finding and map reading) available across the two batteries. This resulted in six tests loading onto a first factor of ‘*Spatial Orientation*’: navigation according to directions, navigation according to landmarks, map reading, route memory and two tests originally part of the object-based battery, Elithorne maze and mazes. The second factor of ‘*Object Manipulation’* included the eight remaining tests part of the of object-based battery along with the scanning and perspective-taking measures included the spatial orientation battery (Figure S2). However, this model did not provide a good fit for the data (supplementary Table S5).

The last model we examined was based on the structure of the correlations observed between the 16 spatial tests (Figure 4), which clustered into three main components. Consequently, this fourth model included three factors representing individual differences in: (1) *Object Manipulation*, (2) *Navigation* and (3) *Visualization* abilities (Figure 5b). This model provided a good fit for the data (χ^2^ = 351.870 (101), p < .001, CFI = 0.953, TLI = 0.944, RMSEA = 0.041, SMRS = 0.041). However, the three factors were strongly correlated (*r* ranging from .73 to .95). Based on these strong correlations, we re-specified the model as a hierarchically-structured model of spatial skills: The 16 tests of spatial skills clustered onto three separate abilities (object manipulation, navigation and visualization), which in turn loaded onto a common factor of *Spatial Ability* (Figure 6).

**Figure 6.**
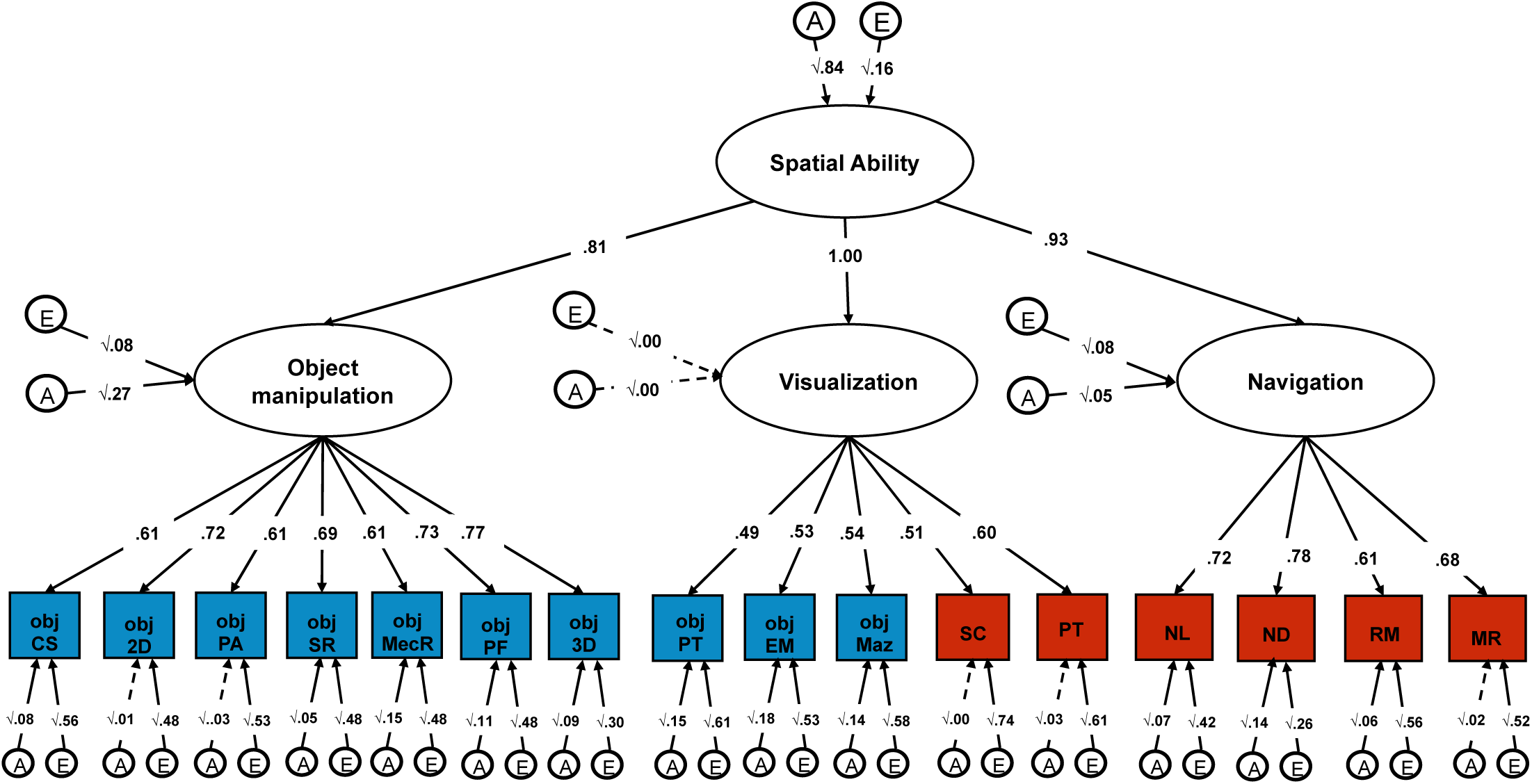
Genetic and environmental variance characterizing the hierarchical structure of spatial ability. Within each blue rectangle are the ten tests that were included in the object-based spatial battery, while shaded in red are the six tests part of the spatial orientation battery set in a naturalistic virtual environment. Both the phenotypic (χ^2^ = 351.870 (101), p < .001, CFI = 0.953, TLI = 0.944, RMSEA = 0.041, SMRS = 0.041) and genetic (χ^2^ = 1681.128 (1040), *p* = 0.0000; CFI = 0.941; TLI = 0.944; RMSEA = 0.026; SRMR = 0.056) model provided good fit for the data.

This hierarchical characterization of spatial skills describes the complexity of the structure of individual differences in spatial abilities, while highlighting the strong interconnection between all abilities at a higher level of analysis. The higher order factor of spatial ability accounted for a large portion of individual differences in the navigation (R^2^ = .791), object manipulation (R^2^ = .689) and visualization (R^2^ = 1.00) factors. We adopted this hierarchical characterization of individual differences in spatial skills in subsequent analyses.

### A common genetic network underlies performance in all spatial tests

We used multivariate twin analysis to analyse the genetic and environmental origins of the hierarchical structure of spatial abilities. First, we found that a model decomposing variation in spatial abilities into additive genetic (A) and nonshared environmental (E) factors provided a good fit for the data (χ^2^ = 1681.128 (1040), p < 0.0005, CFI = 0.941, TLI = 0.944, RMSEA = 0.026, SMRS = 0.056). That is, there was no evidence that shared environmental variance, which encompasses those experiences that make children growing up in the same family more similar to one another beyond their genetic similarity, played a meaningful role in accounting for individual differences in spatial skills.

This hierarchical AE model (Figure 6) showed that spatial skills clustered together largely due to shared genetic variance. The common spatial ability factor was in fact highly heritable (84%) and subsumed 67% of the genetic variance in object manipulation. This is calculated, based on path tracing, as the standardized squared genetic variance in the general factor of spatial ability (.84) multiplied by twice the path estimate for object manipulation (.81) divided by the total genetic variance (.84 *.81^2+.27), resulting in (.84*.81^2)/(.84*.81^2+.27). The common factor of spatial ability accounted for 93% of the genetic variance in the navigation factor and for the entirety of the genetic variance in the visualization factor (see Supplementary Table S6 for the full model including 95% confidence intervals). Nonshared environmental variance accounted for a much smaller proportion of individual differences in the common spatial ability factor (16%).

### General cognitive ability (g) only partly accounts for the genetic clustering of spatial skills

It is well established that cognitive skills correlate with each other, and that a substantial portion of variation in different abilities can be accounted for by a general factor of cognitive ability (*g*), both at the observed and genetic level.^17,35,36^ We applied a Cholesky decomposition (Method) to examine to what extent the genetic and environmental variance in spatial ability could be captured by *g*. The Cholesky approach, similar to hierarchical regression, parses the genetic and environmental variation in each trait into that which is accounted for by traits that have previously been entered into the model and the variance which is unique to a newly entered trait. We applied this method to examine the extent to which the clustering of spatial tests into a common factor of spatial ability could be accounted for by the broader *g* factor. The results presented in Figure 7 (see Figure S3 for the full model) showed that *g* accounted for 55% of the genetic variance in the second-order common spatial ability factor. In other words, 45% of the genetic variance in spatial ability was independent of *g*.

**Figure 7.**
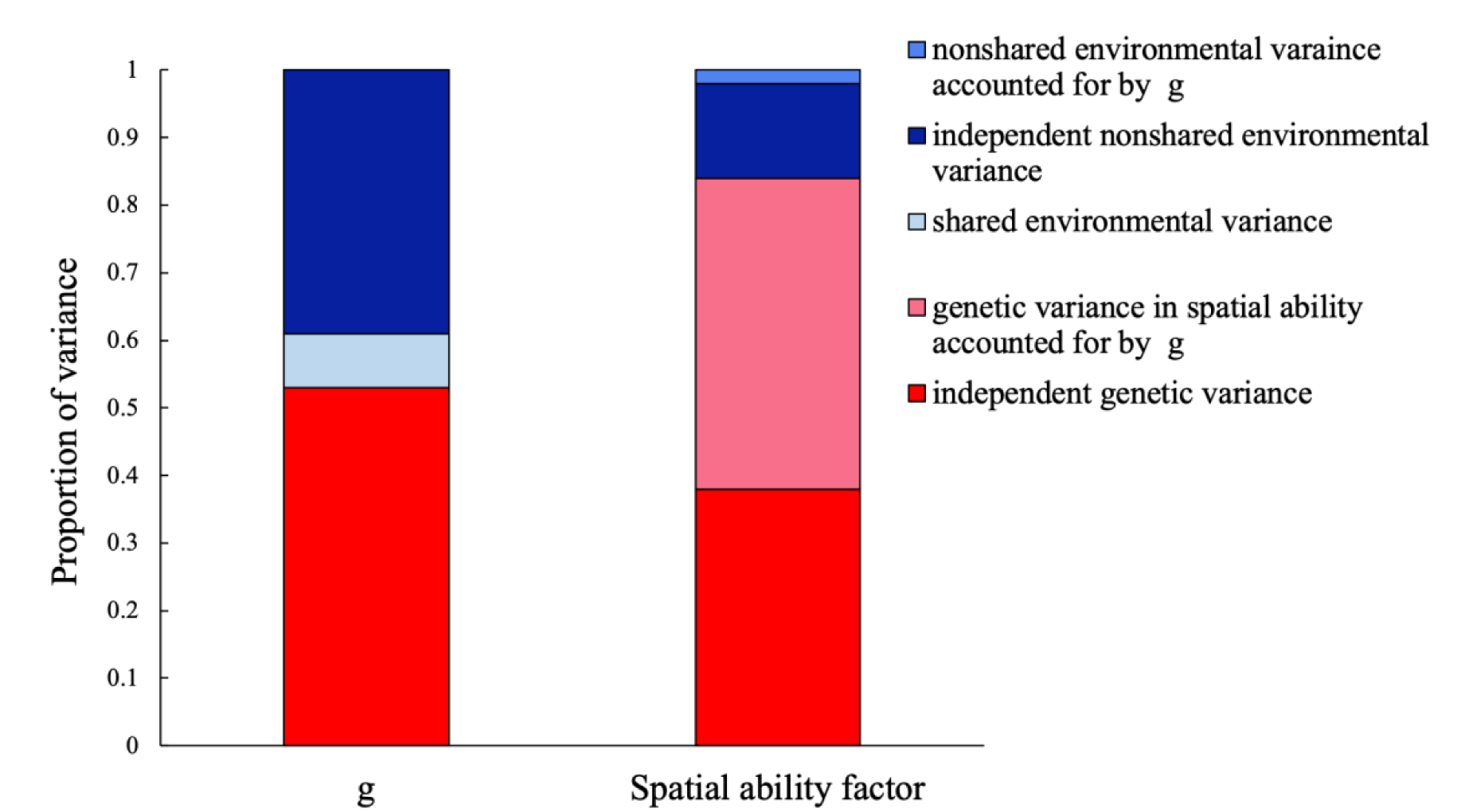
Genetic and environmental variance in a developmental stable measure of *g* and in the common spatial ability factor. For the common spatial ability factor the bar is divided into the genetic and environmental contributions independent of *g* and those that are accounted for by the genetic and environmental variance in *g.* Results are from a Cholesky decomposition (see Supplementary Figure S3 for the full model).

When we accounted for *g* at different levels in the models (Supplementary Figures S4 to S9), results remained consistent with the existence of a general genetic network of spatial skills that covaries independently of *g*.

## DISCUSSION

The current study provides new knowledge on the structure and nature of spatial ability, which addresses three outstanding issues in the field of spatial cognition. First, we examined the structure of spatial orientation abilities, measured with a novel gamified battery set in a virtual environment that included a broad range of measures tapping putatively different aspects of spatial orientation ability. Second, we explored the structure of the associations between spatial orientation skills and object-based spatial tests, a topic that remains mostly unexplored in the cognitive psychology literature and is characterized by strong, contrasting theoretical views. ^7,15,22^ Third, we investigated the extent to which an index of the developmentally stable component of *g* accounted for the shared variance observed across spatial skills. Across these three broad aims, we leveraged the genetically informative quality of our twin sample to address parallel questions related to the genetic and environmental structure of spatial ability and of its association with *g*. At every level of analysis our results highlighted communalities rather than differences across tests of spatial ability, largely supporting a unitary structure of spatial cognition.

Support for the unitary structure of spatial cognition first emerged from phenotypic analyses of our battery of spatial orientation tasks. This finding of a strong general component of variation was remarkable given the breath of spatial orientation skills covered by our newly developed battery. In fact, the development of this innovative, gamified, battery set in a virtual environment was guided by a careful process of literature review aimed at covering all the main domains of spatial orientation described in the existing literature. This resulted in six broad domains that ranged from navigation according to directions and large-scale perspective taking, which, based on Newcombe and Shipley’s (2015) taxonomy, could be categorized as extrinsic-dynamic spatial abilities, to route memory and large-scale scanning, which, based on the same taxonomy, could be described as extrinsic-static spatial abilities^7^. Although extrinsic-static and extrinsic-dynamic abilities have been proposed to be separate skills,^7^ and a meta-analysis of the effects of training spatial ability partly supported this distinction for a few selected tests, ^37^ our results contradict this largely theoretical taxonomy.

We found support for a unitary structure of spatial orientation skills not only at an observed (phenotypic) level, but also in terms of the genetic and environmental factors supporting spatial orientation skills. We found that a common factor of ‘navigation ability’ that was 64% heritable and captured between 66% and 100% of the heritability of the six individual tests of spatial orientation, and to a lesser extent their nonshared environmental variance (between 10% and 36%). This suggests that, to the extent that measures of spatial orientation covary, they do so largely due to their shared genetic variance. These results push our knowledge of the nature of spatial orientation skills further, providing support for a unitary structure of spatial orientation skills at the genetic level.

Further support for a unitary structure of spatial cognition emerged when we considered an even greater breadth of spatial tests, including, in addition to our six measures of spatial orientation, ten psychometric tests of object-based spatial skills, administered in the same sample as part of another gamified spatial battery. These sixteen tests of spatial skills were specifically selected to cover all the main areas of spatial cognition identified in extant literature, making the current work, to our knowledge, the most comprehensive investigation of spatial abilities to date. We approached the examination of the structure of associations between such a broad umbrella of spatial measures by moving through increasing levels of complexity.

A simple unitary account of spatial ability, represented by a general factor common to all measures, was found not to provide an accurate description of the foundations of spatial skills. At a first glance, the results could have been interpreted as supporting the existence of three factors of spatial ability. These three factors described individual differences in navigation, object-based abilities and visualization. Existing taxonomies of spatial ability,^7^ differentiate not only between static and dynamic spatial skills, but also between intrinsic and extrinsic abilities. Consistent with this account we observed a partial differentiation between object-based spatial tests, such as mental rotation, that are largely concerned with the intrinsic properties of objects, and visualization tests, such as perspective taking and scanning, which are largely concerned with extrinsic relations among objects.^7,38^ However, the very strong correlations, from .73 to .95, observed between the object-based, navigation and visualization factors contradicted this putative distinction, and opened the possibility that a coherent, underlying set of abilities held these three factors together.

Factor analytic evidence supported this hierarchical account of spatial cognition: All sixteen tests were found to load onto three factors (navigation ability, object-based ability and visualization ability), which in turn loaded strongly onto a common factor of spatial ability. A hierarchical structure, which highlights both communalities and differences between cognitive tests, has also been found to provide the most accurate characterization in other domains of cognition, most notably executive functions.^39–42^ Also consistent with what has been observed for individual differences in executive functions, we found that genetic factors were largely shared across all tests of spatial abilities. These results point to the existence of a common genetic network at the basis of individual differences in spatial ability, therefore providing additional support for a unitary account of spatial cognition.

A further line of evidence supporting the existence of a unitary account of spatial cognition was provided by our analyses examining the role of *g* in the clustering of spatial ability at the genetic and environmental levels. We found that individual differences in *g* correlated moderately with all individual tests of spatial skills and substantially with the common spatial ability factor. However, nearly half of the substantial genetic variance in spatial ability was found to be independent of the genetic variance in *g*, measured aggregating multiple cognitive tests over development. Taken together, our results indicate that spatial skills cluster together phenotypically and genetically beyond the simple fact that they are all tests reflecting a general, developmentally stable, capacity for planning, thinking abstractly and solving problems, all skills that are indexed by *g.*^35^ It should be noted that, since the genetic and environmental components of cognitive abilities have differential longitudinal stabilities, aggregating across waves might have resulted in ‘cancelling out’ environmental variance that is specific to each developmental stage, in favour of aggregating stable genetic variance in *g* over development.^36^

In summary, our current work provides a threefold line of support for the unitary nature of spatial cognition, partly independent of other measures of cognitive skills. Interestingly, this unitary account of abilities is at odds with individuals’ perceptions of their own ability and feelings towards spatial activities. In our previous work examining the structure of spatial and mathematics anxiety, we found evidence for a separation between the anxiety people feel towards spatial navigation and the anxiety towards object-based skills, such as completing difficult jigsaw puzzles and building flat-pack furniture from instructions.^43^ This observed difference in perceptions and feelings towards different spatial activities might contribute to explaining why ideas, theories and taxonomies of spatial cognition have mostly favoured a multifaceted account of spatial skills.

Although our study provides the most comprehensive investigation of the structure of spatial ability to date in a large sample and addresses several outstanding research questions concerning spatial cognition, it was limited by the technology available to us at the time. Although we developed a new gamified battery set in a virtual environment to reliably examine individual differences in spatial orientation skills, it is possible that assessment in a computer-simulated environment might not be able to capture individual differences in spatial orientation and navigation as well as does assessment in real life settings. It has been proposed that spatial orientation in computer-simulated environments might reflect an allocentric (object-to-object) approximation of the abilities involved in egocentric (self-to-object) real-life spatial orientation.^44^ However, studies that have examined the reliability of measuring navigation skills in virtual reality, as compared to real life settings, have found good concordance between the two.^13^ While we leveraged the newest technological developments to create a realistic virtual environment to host our gamified test, future studies might explore navigation in virtual reality applying even more immersive tools such as, for example, head-mounted displays (e.g. oculus technology).

Our finding of a unitary structure of spatial cognition across sixteen diverse tests of spatial skills, is likely to inform several disciplines beyond cognitive psychology. Investigations on the nature and structure of spatial ability have concerned researchers in a wide range of scientific disciplines, from evolutionary biology, to neuroscience, ecology and molecular genetics. Our evidence for a largely unitary phenotypic and genetic network supporting individual differences in spatial cognition can serve as a basis for future research on the nature of spatial ability across all these disciplines and will likely provide a shift in our consideration of the architecture of human cognitive abilities. These findings are also likely to inform educational practices and interventions, particularly the development of programs aimed at advancing STEM learning through training spatial skills.^45^

## METHODS

### Sample

Participants were part of the Twins Early Development Study (TEDS), a longitudinal study of twins born in the United Kingdom between 1994 and 1996. The families in TEDS are representative of the British population in their socio-economic distribution, ethnicity and parental occupation. See Rimfeld et al. for additional information on the TEDS sample.^47^ The present study focuses on data collected in a sample of 2,660 TEDS twins aged 19-22 (*M* = 21.23, *SD* = 0.53, *range* = 2.29). TEDS twins completed two online batteries assessing multiple aspects of spatial abilities. One was set in a virtual environment and assessed six aspects of large-scale spatial navigation and orientation skills. 2,660 twins (356 MZ males, 338 MZ females, 650 DZ males, 520 DZ females and 796 opposite sex) took part in this battery (868 complete twin pairs). 74.3% of participants who completed the spatial orientation battery (*N* = 1978, 740 complete pairs) also completed an online battery of tests developed to assess ten aspects of object-based spatial abilities. At least five months passed between the administration of the two batteries. The object-based spatial battery was administered first, starting from May 2015, while the data collection for the spatial orientation battery started in September 2015.

### Measures

#### Spatial orientation battery

Putatively different facets of spatial orientation skills were assessed through a novel gamified battery called ‘Spatial Spy’. Participants were invited to solve a mystery by collecting clues while orienting and navigating around the streets of a virtual environment (Figure 8). The online battery was developed in Unity (https://unity3d.com) by ETT Solutions. After a comprehensive literature review, we identified four core aspects of spatial orientation and navigation skills: 1) navigating when reading a map; 2) navigating based on a previously memorized map or route; 3) navigating following directions (e.g. cardinal points), and 4) navigating using reference landmarks. In addition to these four abilities, the spatial orientation battery included two tests based on paradigms that have been frequently used in the object manipulation spatial literature: perspective-taking and scanning. Two research aims motivated the decision to include these two tests in the battery. First, we aimed to explore how perspective taking and scanning measured within a large-scale spatial framework (i.e. within a more naturalistic context approximating a virtual environment) related to the same abilities assessed within a smaller-scale, object manipulation framework (i.e. psychometric tests collected as part of another online battery). Secondly, due to the innovative and experimental nature of the spatial orientation battery, we included measures of scanning and perspective taking, for which we had corresponding data from more traditional psychometric tests, in order to explore the external validity of assessing spatial skills within this new virtual environment. The measures included in this spatial orientation battery are described in detail below. The statistical properties (distribution characteristics and test-retest reliability) of each measure were assessed through two pilot studies.

**Figure 8.**
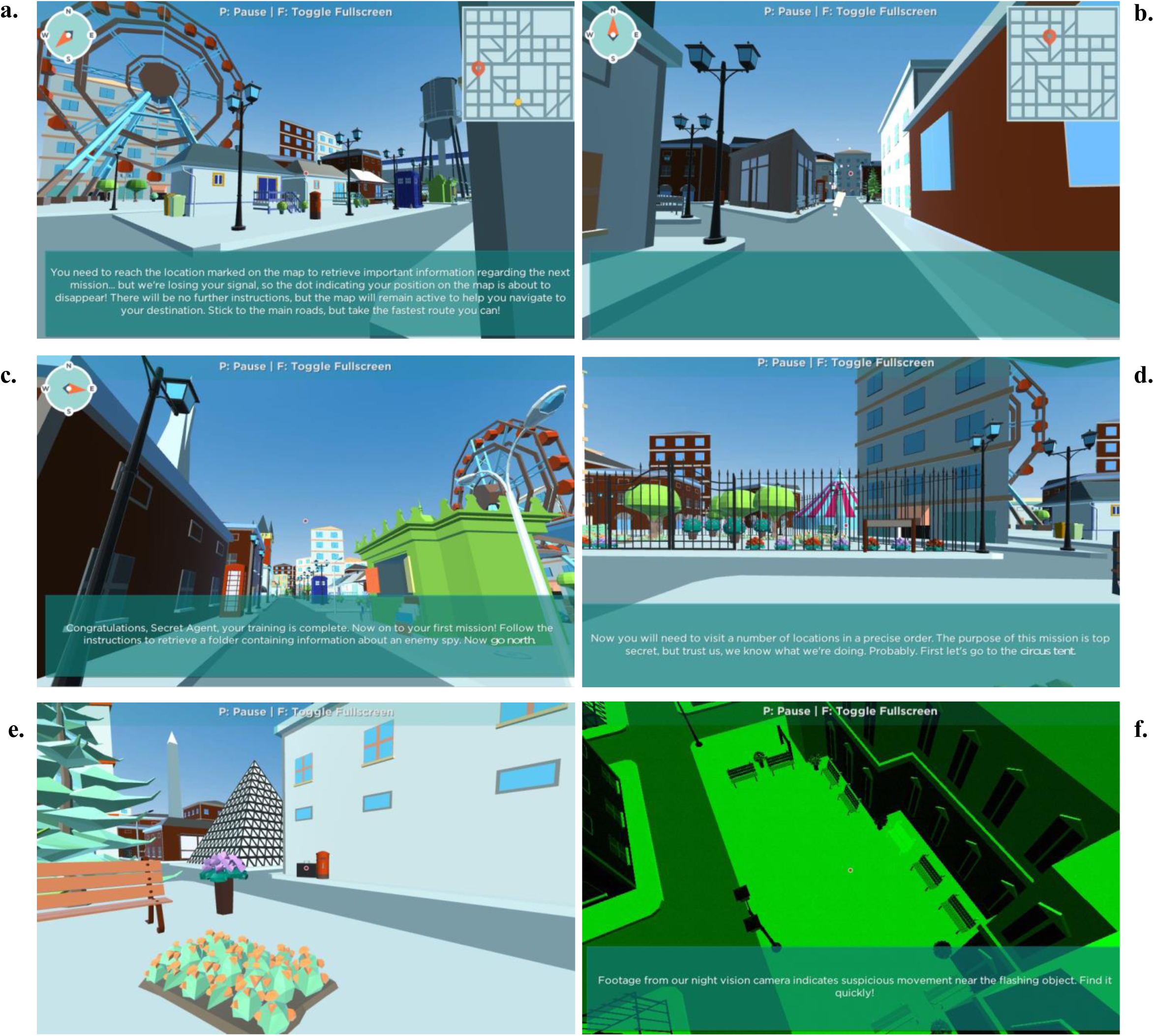
The virtual city where the spatial orientation battery takes place and examples of the six tasks included in the gamified battery: (a) map reading (b) route memorizing; (c) navigation based on directions; (d) navigation based on reference landmarks; (e) scanning; (f) perspective taking.

The final battery started with a training session that helped participants become acquainted with using the cursor or mouse for navigating around the virtual environment, as well as with the requirements and mechanics of each of the six tests. The battery was administered online, with participants taking the tests in web browsers on their own desktop or laptop computers, using a mouse or trackpad to ‘look’ around the virtual environment, and the keyboard to move. The battery took between 35 and 60 minutes (median time 43 minutes) to complete and participants could pause at any time by pressing the key ‘P’ on their keyboard and could resume the game at any given time. Prior to the testing session, participants were provided with practice trials for every test. A two-minute video providing examples of how each subtest was implemented within the Spatial Spy virtual environment is available at the following link https://www.youtube.com/watch?v=wHj0-19rbiI. Following is a detailed description of each test included in the spatial orientation battery. Test-retest reliability for each measure was calculated as part of a pilot study including a sample of 100 participants who completed the battery twice over the space of two months.

##### Map Reading

(Figure 8a), assessed individual differences in the ability to efficiently read a map to travel from one location to another. Once a map had appeared on the top-right corner of the screen, a flashing yellow dot on the map indicated participants’ starting location (A), while a red pointer designated the end-point location on the map (B). Participants were instructed to get from A to B by finding the fastest route and notified that they had 1 minute to complete their mission. If participants could not reach their destination within 60 seconds, they were ‘teleported’ back to the initial location and allowed a second opportunity to complete the task. The ability was assessed though five non-consecutive iterations of increasing difficulty. Each iteration was allocated a score of 2 if participants had successfully travelled from A to B through the quickest (most direct) route, a score of 1 if participants had successful completed the mission but had not selected the fastest route, and a score of 0 if participants had failed to complete the mission. This created a final maximum score of 10. The final score was calculated by combining this accuracy score with participants’ reaction time (time taken to successfully complete the mission), equally weighted. The test showed good test-retest reliability (*r* = .69, p< .001) and distribution (Figure 1).

##### Memorizing a Route

(Figure 8b), assessed individual differences in the ability to travel from one location to another by remembering the content of a map. As for the map reading condition, a map appeared on the top-right corner of the screen, with a flashing yellow dot indicating participant’s starting location (A), and a red pointer designating the end-point location (B). However, the route memorizing test asked participants to memorise the content of the map before the map disappeared from the screen. Participants were given 20 seconds to memorize the map and plan the route before travelling from A to B and were allowed 120 seconds to reach the target location. The number of increasingly difficult iterations, procedure and scoring were the same as those for the previously described map-reading without memory task. Test-retest reliability was good (*r* = .60, p< .001), and distribution (Figure 1).

##### Navigation according to directions

(Figure 8c) assessed participants’ skills in navigating around a virtual environment following instructions based on directions. At the start of the task, participants were ‘teleported’ to one location of the virtual environment and given instructions to navigate around the virtual city in terms of compass points (north, south, east and west). The test included 5 non-consecutive iterations of increasing difficulty and each iteration comprised 4-6 tasks. Each task that was solved correctly was assigned a score of 1. Participants were allowed a maximum of three attempts to respond correctly to each task and consequently proceed to the next set of instructions. After three consecutive failed attempts, the iteration was discontinued and the remaining tasks in that iteration (if any) were assigned a score of 0. Each iteration had a time limit of 180 seconds, if the time limit expired before participants had completed all the tasks, the remaining tasks for that iteration were discontinued and assigned a score of 0. There was no progress bar or timer on screen to help participants keep track of time; however, “hurry up” prompts appeared on screen as the time limit approached. At the end of each iteration (either successfully completed or discontinued) participants were teleported to another part of the virtual environment to complete the following iteration. For the first two iterations the image of a compass providing cardinal directions was available on the top-left corner of the screen, but the compass was not available for the last three iterations, making them more difficult to complete. Examples of instructions were: ‘*Now turn east*’ and ‘*You are facing southwest. Go north and immediately turn west*’. The final score was calculated by combining the accuracy score with participants’ reaction time (time taken to successfully complete each iteration), equally weighted. The test showed excellent test-retest reliability (*r* = .89, p< .001) and distribution of the scores (see Figure 1).

##### Navigating based on reference landmarks

(Figure 8d) measured the ability to navigate following instructions based on the descriptive features of the destination or other nearby landmarks. The test included 5 non-consecutive iterations each comprising 4 or 5 tasks. Each task lasted for a maximum of 60 seconds, so participants had 60 seconds to reach a certain landmark within the virtual environment. If the time limit expired before participants had reached the required landmark, they were discontinued, teleported to the landmark in question, and were able to proceed to the next task. Each task solved correctly, meaning that participants were able to reach the described landmark within the time limit, was assigned a score of 1, while for each trial when participants were not able to reach the location in 60 seconds, they were assigned a score of 0. Neither a map nor a compass was provided to help participants navigate around the environment. Examples of instructions are: ‘*Now reach the tall white pyramid skyscraper*’, and ‘*The message instructs you to go to the park near the old clock tower*’. The target landmark was visible at the start of the session, but it was not always in plain sight as participants were navigating throughout the city to reach the target landmark. The final score was calculated by combining this accuracy score with participants’ reaction time (time taken to successfully complete each iteration), equally weighted. The test showed excellent test-retest reliability (*r* = .80, p< .001) and distribution of the scores (see Figure 3a).

##### Large-scale scanning ability

(Figure 8e) measured participants’ ability to quickly process visual information and identify a target object, a black briefcase, located somewhere nearby within the virtual city. The target object remained the same across the five non-consecutive iterations of increasing difficulty. When looking for the target, participants’ perspective could be rotated freely in any direction, but could not be moved vertically or horizontally around the virtual environment. Participants could identify the target object by clicking on the mouse or trackpad within 60-seconds. Within the time limit, participants were allowed four attempts to correctly spot the target object and, as for all other tasks, they were encouraged to do it as quickly as possible. Feedback was provided after each attempt, and as soon as participants had identified the target object correctly, they were ‘teleported’ to the next task. It was not possible to pause half-way through the 60-second iteration. The final score was calculated by combining this accuracy score with participants’ reaction time (time taken to successfully complete each iteration). The test showed excellent test-retest reliability (*r* = .80, p< .001) and wide distribution of the scores (see Figure 1).

##### Large-scale perspective taking

(Figure 8f) measured participants’ ability to identify objects from a different perspective in large-scale ‘naturalistic’ settings. The test comprised five iterations of increasing difficulty that followed the same test rules. Each iteration started with a CCTV-like image showing an aerial shot of a location within the virtual world, and within this location one target object was depicted flashing on screen for ten seconds. During this initial stimulus presentation, participants could not move within the virtual environment, so all participants were exposed to the same image of the flashing target object. After the ten seconds had elapsed, the CCTV image disappeared and participants were teleported back to the target location within the virtual environment, which shifted their perspective shifted back to ground level; they were then instructed to identify the target object as quickly as possible. When looking for the target object (the one that was flashing when presented from the CCTV perspective), participants’ perspective could be freely rotated but could not be moved vertically or horizontally around the virtual environment. Participants could identify the target object by clicking on it with their mouse or trackpad within 60-seconds. Within the time limit, participants were allowed four attempts to correctly spot the target object and they were encouraged to do it as quickly as possible. A message would appear on the screen after each attempt (either ‘Yes’ or ‘Try again’) to provide participants with feedback on their performance, and each iteration terminated either after a successful attempt, or after participants had used up their four attempts, or if they timed out. A ‘Hurry up’ message was displayed on the screen a few seconds before the time for each iteration elapsed. The test showed good distribution (Figure 1) and test-retest reliability (*r* = .67, p< .001).

#### Object Manipulation

Object manipulation was tested using an online, gamified, battery called ‘The King’s Challenge’^18^. This test battery measures the major putative dimensions of spatial ability, and is comprised of 10 tests: 1) a mazes task (searching for a way through a 2D maze in a speeded task); 2) 2D drawing (sketching a 2D layout of a 3D object from a specified viewpoint); 3) Elithorn mazes (joining together as many dots as possible from an array); 4) pattern assembly (visually combining pieces of objects together to make a whole); 5) mechanical reasoning (multiple-choice naïve physics questions); 6) paper folding (visualizing where the holes are situated after a piece of paper is folded and a hole is punched through it); 7) 3D drawing (sketching a 3D drawing from a 2D diagram); 8) mental rotation (mentally rotating objects); 9) perspective-taking (visualizing objects from a different perspective), and 10) cross-sections (visualizing cross-sections of objects). The development of the battery is described in detail elsewhere.^18^ A brief demonstration of the battery can be accessed here: https://www.youtube.com/watch?v=awnfeiAPmQc

#### General cognitive ability (g) over development

General cognitive ability (*g*; intelligence) was assessed in TEDS at ages 7, 9, 10, 12, 14, and 16. For the present analyses we created a longitudinal composite measure of *g* as a mean of these six assessments. See Supplementary Methods for a more detailed description of *g* measures.

### Analytic Strategies

The R package ‘psych’^48^ was used to obtain descriptive statistics and correlations, and the R package ‘ggplot2’^49^ was used for data visualization purposes. For all phenotypic analyses, one twin was selected randomly from each pair to ensure independence of data. Similar results were obtained when the analyses were conducted on the second half of the sample (see supplementary Figures S10, S11 and S12). Structural Equation modelling (SEM) was conducted in Mplus version 8^50^, and Full Information Maximum Likelihood (FIML) was applied to account for missingness in the data.

#### Confirmatory Factor Analyses

Confirmatory Factor Analysis (CFA) is a data reduction technique whereby latent factors are constructed from observed (measured) indicators based on a pre-imposed structure which is hypothesized to underlie the data. CFA is, in most instances, theory-driven and allows for testing hypothesis on the associations between variables and their underlying latent constructs. Alternative theoretical models were compared examining multiple model fit indices. Model fit indices include: a) the Chi-square test, which indicates the correspondence between the expected and the observed covariance matrices, a chi-square value close to zero indicates greater correspondence between them; b) the Comparative Fit Index (CFI) is an incremental fit index that is based on the non-centrality measure. The CFI ranges from 0 to 1.00, with values closer to 1.00 indicating better fit (acceptable values > .90); c) the Root Mean Square Error of Approximation (RMSEA) is related to residual in the model. RMSEA values range from to 1 with a smaller RMSEA value indicating better model fit. Acceptable model fit is indicated by an RMSEA value of 0.08 or less.^51^

#### Genetic Analyses: Univariate and Multivariate Twin modelling

The twin method allows for the decomposition of individual differences in a trait into genetic and environmental sources of variance by capitalizing on the genetic relatedness between monozygotic twins (MZ), who share 100% of their genetic makeup, and dizygotic twins (DZ), who share on average 50% of the genes that differ between individuals. The method is further grounded in the assumption that both types of twins who are raised in the same family share their rearing environments to approximately the same extent, ^52^ By comparing how similar MZ and DZ twins are for a given trait (intraclass correlations), it is possible to estimate the relative contribution of genetic factors and environments to variation in that trait. Heritability, the amount of variance in a trait that can be attributed to genetic variance (A), can be roughly estimated as double the difference between the MZ and DZ twin intraclass correlations.^53^ The ACE model further partitions the variance into shared environment (C), which describes the extent to which twins raised in the same family resemble each other beyond their shared genetic variance, and non-shared environment (E), which describes environmental variance that does not contribute to similarities between twin pairs (and also includes measurement error). It also provides confidence intervals for all estimates.

When data are available from opposite-sex and same-sex DZ twin pairs, the standard univariate ACE model can be extended to a sex-limitation model to test for the differences in the etiologies of sex differences by comparing five sex and zygosity groups: MZ females, DZ females, MZ males, DZ females and DZ opposite-sex twin pairs. ^32^ This method allows for estimating quantitative and qualitative sex differences (i.e., the same factors affecting males and females to a different extent). Differences in the magnitude of ACE estimates for males and females are referred to as quantitative sex differences; qualitative sex differences indicate whether different genetic or environmental factors influence males and females. The sex limitation model is described in detail elsewhere.^54^ Here we conducted sex-limitation model-fitting by fitting a series of nested models and then testing the relative drop of the fit between the models when the parameters for the sexes are forced to be equal.^55^

The twin method can also be extended to the exploration of the covariance between two or more traits (**multivariate genetic analysis**). Multivariate genetic analysis allows for the decomposition of the covariance between multiple traits into genetic and environmental sources of variance, by modelling the cross-twin cross-trait covariances. Cross-twin cross-trait covariances describe the association between two variables, with twin 1’s score on variable 1 correlated with twin 2’s score on variable 2, which are calculated separately for MZ and DZ twins. The examination of shared variance between traits can be further extended to test the etiology of the variance that is common between traits and of the residual variance that is specific to individual traits. Here we used the **common pathway model** which is is a multivariate genetic model in which the variance common to all measures included in the analysis can be reduced to a common latent factor, for which the A, C and E components are estimated. As well estimating the etiology of the common latent factor, the model allows for the estimation of the A, C and E components of the residual variance in each measure that is not captured by the latent construct. ^56^ The common pathway model estimates the extent to which the general factor of spatial ability is explained by A, C and E. The common pathway model is illustrated in Figure 3. Based on factor analytic evidence, the common pathway model can be extended to include multiple common factors and, consequently to the examination of the genetic and environmental associations between the multiple latent factors. This extension of the common pathway model is presented in Figure 6.

## Acknowledgements

We gratefully acknowledge the ongoing contribution of the participants in the Twins Early Development Study (TEDS) and their families. TEDS is supported by a program grant to R.P. from the UK Medical Research Council [MR/M021475/1 and previously G0901245], with additional support from the US National Institutes of Health [HD044454; HD059215]. R.P. is supported by a Medical Research Council Research Professorship award [G19/2] and a European Research Council Advanced Investigator award [295366]. E.M.T. and M.M. are Faculty Research Associates of the Population Research Center at the University of Texas at Austin, which is supported by a grant, 5-R24-HD042849, from the Eunice Kennedy Shriver National Institute of Child Health and Human Development (NICHD). E.M.T. is also supported by a Jacobs Foundation Research Fellowship and NIH grant R01HD083613. M.M. is partly supported by a David Wechsler Early Career Grant for Innovative Work in Cognition. The funders had no role in study design, data collection and analysis, decision to publish, or preparation of the manuscript.

## Author Contributions

Conceived and designed the experiment: M.M., K.R., N.G.S., K.L.S., M.R., Y.K., V.R., R.P. Analyzed the data: M.M., K.R., N.G.S. Wrote the paper: M.M., K.R., R.P. All authors approved the final draft.

## Supplementary information accompanies this manuscript

### Data Access

the Twins Early Development Study’s data access policy can be found at the following link https://www.teds.ac.uk/researchers/teds-data-access-policy

