## Supplementary Material for "Evidence for a unitary structure of spatial cognition beyond general intelligence"

**Supplementary Methods**

Measures of general cognitive ability (g) over development

**List of Tables**

**Table S1.** **(a)** Descriptive statistics for each spatial test and *g*, randomly selecting one twin out of each pair. **(b)** Descriptive statistics for the other half of the sample.
**Table S2.** Univariate Sex limitation models for individual tests of navigation included in the Spatial Spy battery
**Table S3.** Confirmatory factor analysis and model fit indices across the six tests of spatial orientation
**Table S4.** Common pathway model across the six tests of spatial orientation
**Table S5.** Comparative model fit indices for the phenotypic models including all 16 tests of spatial skills (6 tests of spatial orientation and 10 tests of object-based spatial skills).
**Table S6.** Hierarchical common pathway model examining the genetic and environmental factors characterizing variation in across the 16 tests of spatial abilities (6 tests of spatial orientation and 10 tests of object-based spatial skills).

**List of Figures**

**Figure S1.** Two-factor model of spatial ability separating across the two spatial batteries administered at two different time points and following two different formats (online traditional psychometric assessment vs. virtual environment).
**Figure S2.** Two-factor model of spatial ability separating objects-based and orienting tests combining putatively separate categories of tests administered across the two batteries.
**Figure S3.** Cholesky decomposition exploring the genetic and environmental overlap between *g* and the hierarchical model of spatial ability factor

**Figure S4.** One-factor model of spatial ability including the 16 spatial tests accounting for *g* at the level of the indicator (each test).
**Figure S5.** Two-factor model of spatial ability separating the two batteries accounting for *g* at the level of the indicators (each test).
**Figure S6.** Two-factor model of spatial ability separating objects-based and orienting tests combining putatively different aspects of spatial skills across the two batteries accounting for *g* at the level of the indicators (each test).
**Figure S7.** Three-factor model of spatial ability separating objects-based, navigation and visualization tests across the two batteries accounting for *g*.
**Figure S8.** Hierarchical model including three first-order spatial factors (Navigation, Object-based and Visualization) and a second-order common factor of Spatial ability, accounting for *g* at the level of the indicators (each test).
**Figure S9.** Hierarchical model including three first-order spatial factors (Navigation, Object-based and Visualization) and a second-order common factor of Spatial ability, accounting for *g* at the level of the first-order factors.
**Figure S10.** Correlations between all spatial tests on the other half of the sample.
**Figure S11.** Factor structure of navigation abilities conducted examining the other half of the sample
**Figure S12.** Hierarchical model of spatial abilities conducted in the other half of the sample.

**Supplementary Methods**

***Measures of general cognitive ability (g) over development***

General cognitive ability ( *g* ; intelligence) was assessed in TEDS at ages 7, 9, 10, 12, 14, and 16. For the present analyses we created a longitudinal composite measure of *g* as a mean of these six assessments. At age 7, 'g' was calculated as a mean of conceptual grouping[1], a WISC similarities test[2], a WISC vocabulary test[2], and a WISC picture completion test[2] all collected over telephone testing. At age 9, 'g' was calculated as a mean of a shapes test[3], a WISC vocabulary test[4], a WISC general knowledge task[4], and a puzzle test[3] ; all tests were administered in booklets sent to the twins by post. At age 10, 'g' was calculated as a mean of the Ravens Standard Progressive Matrices[5], a WISC vocabulary[4], WISC picture completion[2], and a WISC general knowledge test[4]; at age 10 and subsequent assessments, all 'g' data were obtained by internet testing. At age 12, 'g' was calculated as a mean of the Ravens Progressive Matrices[5], a WISC picture completion[2], a WISC vocabulary[4], and a WISC general knowledge test[4.] At age 14, 'g' was calculated as a mean of the Raven's Progressive Matrices[5] and a WISC vocabulary[4]. At age 16, 'g' was calculated as a mean of Mill Hill Vocabulary test[6] and Raven's Progressive Matrices[5].

**Tables**

**Table S1. (a)** Descriptive statistics for each spatial test and *g*, randomly selecting one twin out of each pair. **(b)** Descriptive statistics for the other half of the sample.

**(a)**

| | N | mean | sd | min | max | range | skew | kurtosis |
|---|---|---|---|---|---|---|---|---|
| Navigation directions (cardinal points | 1330 | 0.000 | 1.000 | -3.220 | 2.440 | 5.660 | -0.170 | -0.300 |
| Navigation landmarks | 1278 | 0.000 | 1.000 | -4.260 | 1.980 | 6.240 | -1.140 | 1.610 |
| Map reading | 1244 | 0.000 | 1.000 | -4.180 | 1.620 | 5.800 | -1.020 | 1.040 |
| Route memorizing | 1217 | 0.000 | 1.000 | -4.260 | 1.510 | 5.770 | -1.350 | 1.920 |
| Large-scale perspective taking | 1285 | 0.000 | 1.000 | -3.350 | 1.470 | 4.830 | -0.960 | 0.400 |
| Large-scale scanning | 1229 | 0.000 | 1.000 | -4.160 | 1.590 | 5.750 | -1.430 | 2.120 |
| KC cross-section | 927 | 0.000 | 1.000 | -2.200 | 2.310 | 4.510 | -0.180 | -0.760 |
| KC 2d drawing | 920 | 0.000 | 1.000 | -3.480 | 1.540 | 5.020 | -0.710 | -0.020 |
| KC pattern assembly | 896 | 0.000 | 1.000 | -2.260 | 2.620 | 4.870 | -0.370 | -0.520 |
| KC Elithorne maze | 804 | 0.000 | 1.000 | -3.210 | 1.990 | 5.200 | -1.160 | 1.290 |
| KC mechanical reasoning | 914 | 0.000 | 1.000 | -3.060 | 2.940 | 6.000 | -0.180 | -0.050 |
| KC paper folding | 888 | 0.000 | 1.000 | -2.290 | 2.090 | 4.390 | -0.120 | -1.000 |
| KC 3d drawing | 833 | 0.000 | 1.000 | -2.030 | 2.340 | 4.370 | 0.030 | -1.000 |
| KC shapes rotation | 850 | 0.000 | 1.000 | -2.330 | 1.870 | 4.200 | -0.340 | -0.850 |
| KC small-scale perspective taking | 859 | 0.000 | 1.000 | -1.640 | 2.980 | 4.620 | 0.610 | -0.190 |
| KC mazes | 850 | 0.000 | 1.000 | -3.070 | 2.650 | 5.720 | -0.140 | -0.120 |
| General cognitive ability (g) | 1234 | 0.000 | 1.000 | -2.830 | 3.040 | 5.870 | 0.320 | 0.010 |

**Note:** all measures were standardized and residualized for age and sex within the randomly selected half of the sample by means of linear regression

**(b)** Descriptive statistics for the other half of the sample

| | N | mean | sd | min | max | range | skew | kurtosis |
|---|---|---|---|---|---|---|---|---|
| Navigation directions (cardinal points | 1349 | 0.000 | 1.000 | -3.250 | 2.790 | 6.050 | -0.110 | -0.330 |
| Navigation landmarks | 1312 | 0.000 | 1.000 | -4.330 | 2.070 | 6.400 | -0.900 | 0.870 |
| Map reading | 1268 | 0.000 | 1.000 | -4.300 | 1.620 | 5.930 | -1.000 | 1.160 |
| Route memorizing | 1234 | 0.000 | 1.000 | -4.500 | 1.550 | 6.050 | -1.250 | 1.710 |
| Large-scale perspective taking | 1316 | 0.000 | 1.000 | -3.450 | 1.450 | 4.900 | -0.850 | 0.170 |
| Large-scale scanning | 1260 | 0.000 | 1.000 | -3.960 | 1.610 | 5.570 | -1.250 | 1.510 |
| KC cross-section | 939 | 0.000 | 1.000 | -2.250 | 2.330 | 4.580 | -0.260 | -0.710 |
| KC 2d drawing | 932 | 0.000 | 1.000 | -3.230 | 1.610 | 4.850 | -0.790 | 0.180 |
| KC pattern assembly | 911 | 0.000 | 1.000 | -2.420 | 2.130 | 4.550 | -0.440 | -0.580 |
| KC Elithorne maze | 815 | 0.000 | 1.000 | -3.850 | 1.940 | 5.790 | -1.060 | 1.280 |
| KC mechanical reasoning | 921 | 0.000 | 1.000 | -3.080 | 2.590 | 5.680 | -0.100 | -0.230 |
| KC paper folding | 893 | 0.000 | 1.000 | -2.380 | 1.900 | 4.280 | -0.270 | -0.960 |
| KC 3d drawing | 868 | 0.000 | 1.000 | -2.130 | 2.240 | 4.380 | -0.090 | -1.010 |
| KC shapes rotation | 857 | 0.000 | 1.000 | -2.450 | 1.920 | 4.370 | -0.420 | -0.760 |
| KC small-scale perspective taking | 880 | 0.000 | 1.000 | -1.680 | 3.180 | 4.850 | 0.590 | -0.450 |
| KC mazes | 862 | 0.000 | 1.000 | -3.170 | 2.550 | 5.720 | -0.220 | -0.180 |
| General cognitive ability (g) | 1242 | 0.000 | 1.000 | -2.770 | 2.950 | 5.720 | 0.150 | -0.150 |

**Note:** all measures were standardized and residualized for age and sex within the randomly selected half of the sample by means of linear regression

**Table S2.** Sex limitation model fitting sub-model comparisons (significant differences are marked in bold). FullHetACE=full genetic heterogeneity model, rG=Free; HetACE= quantitative heterogeneity model; cFullHetACE=full environmental heterogeneity model, rC=Free; HomACE= homogeneity model (no sex differences at all); ep=estimated parameters; minus2LL= minus 2 log-likelihood; df= degrees of freedom; AIC= Akaike information criterion; diffLL= change in log-likelihood; diffdf= change in degrees of freedom (significant differences are marked in bold).

| | | | | | | | |
|---|---|---|---|---|---|---|---|
| **Navigation ability** | | | | | | | |
| *Qualitative genetic differences:* | | | | | | | |
| | ep | -2LL | df | AIC | diffLL | diffdf | p |
| Model: FullHetACE | 9 | 5863.64 | 2121 | 1621.64 | - | - | - |
| Model: HetACE | 8 | 5863.64 | 2122 | 1619.64 | 0 | 1 | 0.96 |
| *Qualitative environmental differences:* | | | | | | | |
| | ep | -2LL | df | AIC | diffLL | diffdf | p |
| Model: cFullHetACE | 9 | 5870.66 | 2121 | 1628.66 | - | - | - |
| Model: HetACE | 8 | 5863.64 | 2122 | 1619.64 | -7.01 | 1 | 1 |
| Quantitative differences: | | | | | | | |
| | ep | -2LL | df | AIC | diffLL | diffdf | p |
| Model: HetACE | 8 | 5863.64 | 2122 | 1619.64 | - | - | - |
| Model: HomACE | 5 | 5874.35 | 2125 | 1624.35 | 10.71 | 3 | 0.01 |
| **Scanning** | | | | | | | |
| *Qualitative genetic differences:* | | | | | | | |
| | ep | -2LL | df | AIC | diffLL | diffdf | p |
| Model: FullHetACE | 9 | 6992.87 | 2480 | 2032.87 | - | - | - |
| Model: HetACE | 8 | 6992.87 | 2481 | 2030.87 | 0 | 1 | 1 |

*Qualitative environmental differences:*

| | ep | -2LL | df | AIC | diffLL | diffdf | p |
|---|---|---|---|---|---|---|---|
| Model: cFullHetACE | 9 | 7006.02 | 2480 | 2046.02 | - | - | - |
| Model: HetACE | 8 | 6992.87 | 2481 | 2030.87 | -13.15 | 1 | 1 |

Quantitative differences:

| | ep | -2LL | df | AIC | diffLL | diffdf | p |
|---|---|---|---|---|---|---|---|
| Model: HetACE | 8 | 6992.87 | 2481 | 2030.87 | - | - | - |
| Model: HomACE | 5 | 7048.2 | 2484 | 2080.2 | 55.33 | 3 | 0 |

**Perspective taking**

*Qualitative genetic differences:*

| | ep | -2LL | df | AIC | diffLL | diffdf | p |
|---|---|---|---|---|---|---|---|
| Model: FullHetACE | 9 | 7314.24 | 2592 | 2130.24 | - | - | - |
| Model: HetACE | 8 | 7314.77 | 2593 | 2128.77 | 0.53 | 1 | 0.47 |

*Qualitative environmental differences:*

| | ep | -2LL | df | AIC | diffLL | diffdf | p |
|---|---|---|---|---|---|---|---|
| Model: cFullHetACE | 9 | 7336 | 2592 | 2152 | - | - | - |
| Model: HetACE | 8 | 7314.77 | 2593 | 2128.77 | -21.23 | 1 | 1 |

Quantitative differences:

| | ep | -2LL | df | AIC | diffLL | diffdf | p |
|---|---|---|---|---|---|---|---|
| Model: HetACE | 8 | 7314.77 | 2593 | 2128.77 | - | - | - |
| Model: HomACE | 5 | 7328.79 | 2596 | 2136.79 | 14.02 | 3 | 0 |

**Navigation according to landmarks**

*Qualitative genetic differences:*

| | ep | -2LL | df | AIC | diffLL | diffdf | p |
|---|---|---|---|---|---|---|---|
| Model: FullHetACE | 9 | 7216.09 | 2581 | 2054.09 | - | - | - |
| Model: HetACE | 8 | 7216.09 | 2582 | 2052.09 | 0 | 1 | 1 |

*Qualitative environmental differences:*

| | ep | -2LL | df | AIC | diffLL | diffdf | p |
|---|---|---|---|---|---|---|---|
| Model: cFullHetACE | 9 | 7216.09 | 2581 | 2054.09 | - | - | - |
| Model: HetACE | 8 | 7216.09 | 2582 | 2052.09 | 0 | 1 | 1 |

Quantitative differences:

| | ep | -2LL | df | AIC | diffLL | diffdf | p |
|---|---|---|---|---|---|---|---|
| Model: HetACE | 8 | 7216.09 | 2582 | 2052.09 | - | - | - |
| Model: HomACE | 5 | 7248.5 | 2585 | 2078.5 | 32.42 | 3 | 0 |

**Navigation according to directions**

*Qualitative genetic differences:*

| | ep | -2LL | df | AIC | diffLL | diffdf | p |
|---|---|---|---|---|---|---|---|
| Model: FullHetACE | 9 | 7390.91 | 2670 | 2050.91 | - | - | - |
| Model: HetACE | 8 | 7392.27 | 2671 | 2050.27 | 1.36 | 1 | 0.24 |

*Qualitative environmental differences:*

| | ep | -2LL | df | AIC | diffLL | diffdf | p |
|---|---|---|---|---|---|---|---|
| Model: cFullHetACE | 9 | 7390.91 | 2670 | 2050.91 | - | - | - |
| Model: HetACE | 8 | 7392.27 | 2671 | 2050.27 | 1.36 | 1 | 0.24 |

Quantitative differences:

| | ep | -2LL | df | AIC | diffLL | diffdf | p |
|---|---|---|---|---|---|---|---|
| Model: HetACE | 8 | 7392.27 | 2671 | 2050.27 | - | - | - |
| Model: HomACE | 5 | 7398.39 | 2674 | 2050.39 | 6.12 | 3 | 0.11 |

**Map reading**

*Qualitative genetic differences:*

| | ep | -2LL | df | AIC | diffLL | diffdf | p |
|---|---|---|---|---|---|---|---|
| Model: FullHetACE | 9 | 6980.65 | 2503 | 1974.65 | - | - | - |
| Model: HetACE | 8 | 6983.5 | 2504 | 1975.5 | 2.85 | 1 | 0.09 |

*Qualitative environmental differences:*

| | ep | -2LL | df | AIC | diffLL | diffdf | p |
|---|---|---|---|---|---|---|---|
| Model: cFullHetACE | 9 | 6983.5 | 2503 | 1977.5 | - | - | - |
| Model: HetACE | 8 | 6983.5 | 2504 | 1975.5 | 0 | 1 | 1 |

Quantitative differences:

| | ep | -2LL | df | AIC | diffLL | diffdf | p |
|---|---|---|---|---|---|---|---|
| Model: HetACE | 8 | 6983.5 | 2504 | 1975.5 | - | - | - |
| Model: HomACE | 5 | 7079.69 | 2507 | 2065.69 | 96.19 | 3 | 0 |

**Route memorizing**

*Qualitative genetic differences:*

| | ep | -2LL | df | AIC | diffLL | diffdf | p |
|---|---|---|---|---|---|---|---|
| Model: FullHetACE | 9 | 6769.48 | 2442 | 1885.48 | - | - | - |
| Model: HetACE | 8 | 6769.48 | 2443 | 1883.48 | 0 | 1 | 1 |

*Qualitative environmental differences:*

| | ep | -2LL | df | AIC | diffLL | diffdf | p |
|---|---|---|---|---|---|---|---|
| Model: cFullHetACE | 9 | 6769.48 | 2442 | 1885.48 | - | - | - |
| Model: HetACE | 8 | 6769.48 | 2443 | 1883.48 | 0 | 1 | 1 |

Quantitative differences:

| | ep | -2LL | df | AIC | diffLL | diffdf | p |
|---|---|---|---|---|---|---|---|
| Model: HetACE | 8 | 6769.48 | 2443 | 1883.48 | - | - | - |
| Model: HomACE | 5 | 6894.1 | 2446 | 2002.1 | 124.63 | 3 | 0 |

**Table S3.** Confirmatory factor analysis and model fit indices across the six tests of spatial orientation.

| Spatial orientation battery | Factor loadings | S.E. |
|---|---|---|
| Navigation directions (cardinal points) | 0.736 | 0.017 |
| Navigation landmarks | 0.756 | 0.017 |
| Map reading | 0.682 | 0.019 |
| Route memorizing | 0.636 | 0.021 |
| Large-scale perspective taking | 0.582 | 0.022 |
| Large-scale scanning | 0.566 | 0.024 |
| Model fit indices: $\chi^2$ = 69.051(9), p< .00005, CFI = 0.972 TLI = 0.953, RMSEA = 0.071 SRMR. =0.026 | | |

**Table S4.** Common pathway model examining the common and specific genetic (A), shared-environmental (C) and nonshared environmental variance (E) across the six tests of spatial orientation and model fit indices.

| **Variance specific to each test** | | | | **Percentages of A, C and E variance in each test captured by the common Navigation factor** | | |
|---|---|---|---|---|---|---|
| **Measure** | **A** | **C** | **E** | **A** | **C** | **E** |
| Navigation directions | 0.181 (0.136; 0.233) | 0.001 (-0.000; 0.001) | 0.274 (0.228; 0.324) | 66% | 100% | 36% |
| Navigation landmarks | 0.000 (-0.001, 0.001) | 0.038 (0.006; 0.096) | 0.394 (0.340; 0.451) | 88% | 53% | 28% |
| Map reading | 0.035 (0.000; 0.130) | 0.000 (-0.000; 0.000) | 0.506 (0.431; 0.586) | 88% | 100% | 20% |
| Route memorizing | 0.058 (0.005; 0.167) | 0.001 (-0.078; 0.099) | 0.564 (0.478; 0.654) | 80% | 100% | 16% |
| Large-scale Scanning | 0.000 (-0.001; 0.002) | -0.000 (-0.000; 0.000) | 0.705 (0.664; 0.748) | 100% | 100% | 10% |
| Large-scale Perspective-taking | 0.047 (0.003; 0.139) | 0.000 (0.000; 0.000) | 0.597(0.524; 0.675) | 82% | 100% | 15% |
| **Variance common across all tests** | | | | | | |
| Navigation latent factor | 0.634 (0.410; 0.912) | 0.083 (-0.005; 0.430) | 0.278 (0.206; 0.362) | | | |
| Model fit indices: AIC = 39177.976; $\chi^2$ = 269.937 (148), $p$ = 0.0000; CFI = 0.968; TLI = 0.971; RMSEA = 0.030 ; SRMR = 0.049 | | | | | | |

**Note.** All paths are standardized and squared, numbers in parentheses are 95% confidence intervals around the estimates.

**Table S5.** Comparative model fit indices for the phenotypic models including all 16 tests of spatial skills (6 tests of spatial orientation and 10 tests of object-based spatial skills)

| | **Model** | **CFI** | **TLI** | **RMSEA** | **Chi^2** | **SRMR** | **Correlations between latent factors** |
|---|---|---|---|---|---|---|---|
| a | 1 Factor | 0.890 | 0.873 | 0.061 | 692.730 (104), p < .001 | 0.059 | - |
| b | 1 Factor *accounting for g*[1] | 0.894 | 0.862 | 0.067 | 609.795 (104), p < .001 | 0.053 | - |
| c | 2 Factors (Spatial Spy battery and. King's Challenge battery) | 0.958 | 0.951 | 0.037 | 316.000 (103), p < .001 | 0.040 | .741 |
| d | 2 Factors (Spatial Spy battery and. King's Challenge battery) *accounting for g*[1] | 0.961 | 0.949 | 0.041 | 288.468 (103), p < .001 | 0.038 | .659 |
| e | 2 Factors (Object Manipulation and Spatial Orientation) | 0.920 | 0.907 | 0.053 | 529.390 (103), p< .001 | 0.054 | .847 |
| f | 2 Factors (Object Manipulation and Spatial Orientation) *accounting for g*[1] | 0.925 | 0.901 | 0.057 | 461.763 (103), p < .001 | 0.051 | .775 |
| g | 3 Factors (Object Manipulation Navigation and Visualization) | 0.953 | 0.944 | 0.041 | 351.870 (101), p < .001 | 0.041 | Obj with Or = .726<br>Or with Sc = .948<br>Sc with Obj = .858 |
| h | 3 Factors (Object Manipulation Navigation and Visualization) *accounting for g*[1] | 0.957 | 0.942 | 0.043 | 306.307 (101), p < .001 | 0.038 | Obj with Or = .633<br>Or with Sc = .949<br>Sc with Obj = .806 |
| i | 3 Factors (Object Manipulation Navigation and Visualization) and a second order common Spatial Ability factor | 0.953 | 0.944 | 0.041 | 351.870 (101), p < .001 | 0.041 | |

| | | | | | | | |
|---|---|---|---|---|---|---|---|
| j | 3 Factors (Object Manipulation Navigation and Visualization) and a second order common Spatial Ability factor *accounting for g*[1] | 0.957 | 0.942 | 0.043 | 306.307 (101), p < .001 | 0.038 | - |
| k | 3 Factors (Object Manipulation Navigation and Visualization) and a second order common Spatial Ability factor *accounting for g*[2] | 0.951 | 0.941 | 0.044 | 348.789 (114), p < .001 | 0.041 | - |

**1 =** g included in the model at the level of the indicators; 2 = g included in the model at the level of the first order latent factors. Models h and j and models i and are k represent two different ways f specifying the same model, therefore their model fit indices are identical.

**Table S6.** Genetic and environmental variance components for the hierarchical model of spatial abilities

| | Variance specific to each test | | Loading on first order factor | A and E variance captured by the first order factors but not shared with the general spatial ability factor | |
|---|---|---|---|---|---|
| **Measure** | **A** | **E** | | **A** | **E** |
| Navigation from directions (cardinal points) | 0.136 (0.094; 0.186) | 0.255 (0.214; 0.309) | 0.777 (0.755; 0.799) | 0.030 | 0.048 |
| Navigation from landmarks | 0.072 (0.025; 0.145) | 0.419 (0.356; 0.487) | 0.712 (0.685; 0.740) | 0.025 | 0.041 |
| Map reading | 0.016; (-0.016; 0.133) | 0.511 (0.438; 0.588) | 0.687 (0.658; 0.717) | 0.023 | 0.036 |
| Route memorizing | 0.059 (0.007; 0.160) | 0.565 (0.483; 0.654) | 0.612 (0.577; 0.647) | 0.018 | 0.029 |
| Cross-sections | 0.075 (0.024; 0.155) | 0.556 (0.491; 0.625) | 0.606 (0.573; 0.639) | 0.100 | 0.029 |
| 2D drawing | 0.010 (-0.046; 0.173) | 0.470 (0.398; 0.549) | 0.721 (0.694; 0.748) | 0.139 | 0.041 |
| Pattern assembly | 0.028 (-0.000; 0.128) | 0.527 (0.456; 0.602) | 0.666 (0.637; 0.696) | 0.100 | 0.029 |
| Shapes rotation | 0.051 (0.009; 0.125) | 0.476 (0.412; 0.543) | 0.688 (0.658; 0.718) | 0.128 | 0.036 |
| Mechanical reasoning | 0.150 (0.093; 0.221) | 0.473 (0.405; 0.546) | 0.614; (0.579; 0.648) | 0.100 | 0.029 |
| Paper folding | 0.107 (0.059; 0.169) | 0.366 (0.302; 0.429) | 0.726 (0.698; 0.754) | 0.143 | 0.042 |
| 3D drawing | 0.093 (0.046; 0.155) | 0.303 (0.250; 0.362) | 0.777 (0.752; 0.801) | 0.160 | 0.047 |
| Small-scale perspective taking | 0.154 (0.081; 0.250) | 0.602 (0.512; 0.697) | 0.494 (0.453; 0.535) | 0.000 | 0.000 |
| Large-scale scanning | 0.000 (-0.001; 0.002) | 0.741 (0.698; 0.783) | 0.509 (0.468; 0.550) | 0.000 | 0.000 |

| | | | | | |
|---|---|---|---|---|---|
| Large-scale perspective-taking | 0.024 (-0.000; 0.136) | 0.608 (0.538; 0.688) | 0.604 (0.571; 0.636) | 0.000 | 0.000 |
| Elithorne Mazes | 0.176 (0.096; 0.284) | 0.532 (0.438; 0.636) | 0.537 (0.486; 0.604) | 0.000 | 0.000 |
| Mazes | 0.134 (0.067; 0.223) | 0.577 (0.497; 0.664) | 0.537 (0.495; 0.579) | 0.000 | 0.000 |
| Variance captured by the first order factors | | | **Loading on second order factor** | **A and E variance captured by the general spatial ability factor** | |
| Navigation latent factor | 0.051 (0.011; 0.122) | 0.085 (0.041; 0.145) | 0.929 (0.904; 0.954) | 0.726 | 0.138 |
| Object-based latent factor | 0.265 (0.198; 0.339) | 0.075 (0.028; 0.145) | 0.812 (0.776; 0.848) | 0.551 | 0.104 |
| Visualization latent factor | 0.000 (0.000; 0.000) | 0.000 (0.000; 0.000) | 1.00 (1.00; 1.000) | 0.837 | 0.163 |
| **Variance common to all tests captured by the second order Spatial Ability factor** | | | | | |
| **Common factor of Spatial ability** | 0.837 (0.779; 0.894) | 0.163 (0.110; 0.225) | - | - | - |
| Model fit indices: AIC = 82828.772; $\chi^2$ = 1681.128 (1040), *p* = 0.0000; CFI = 0.941; TLI = 0.944; RMSEA = 0.026 ; SRMR = 0.056 | | | | | |

### Figures

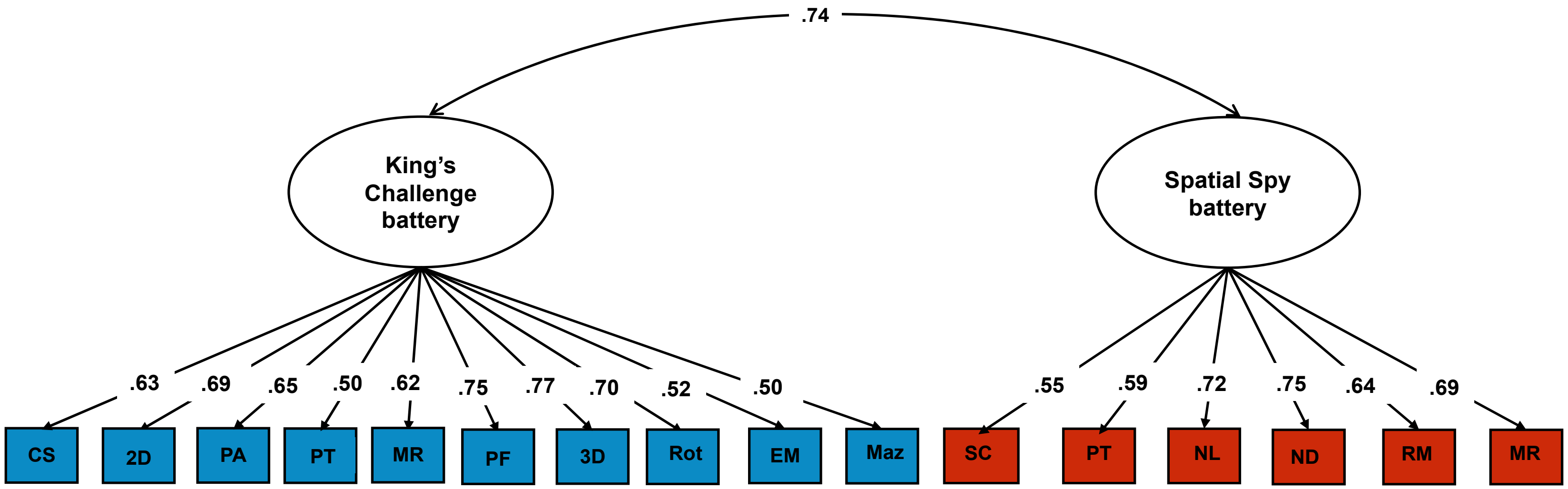


**Figure S1.** Two-factor model of spatial ability separating across the two spatial batteries administered at two different time points and following two different formats (online traditional psychometric assessment – The King's Challenge battery – vs. virtual environment – The Spatial Spy battery). ND = navigation based on directions, NL = navigation based on landmarks, MR = map reading, RM = route memory, PT = perspective taking, SC = scanning, CS = cross-section, 2d = 2d drawing, PA = pattern assembly, EM =Elithorn Mazes, MR = Mechanical Reasoning, PF = paper folding, 3d = 3d drawing, Rot = mental rotation, PT = perspective taking, Maz = mazes; Model fit indices are reported in Table S5.

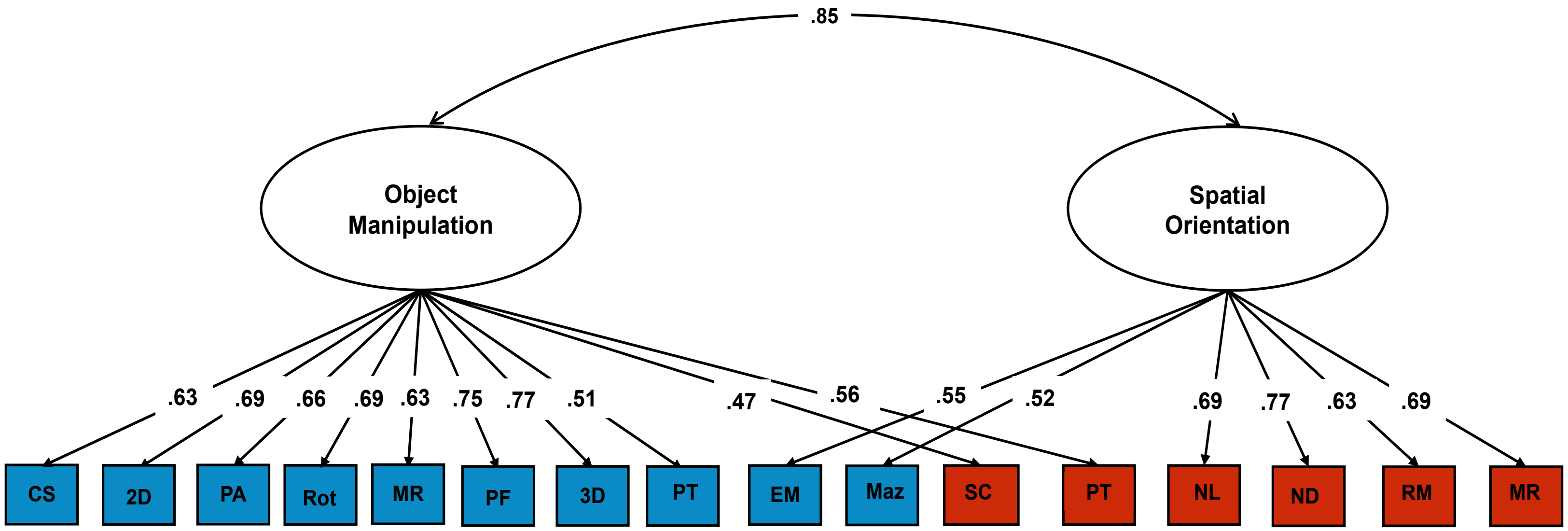


**Figure S2.** Two-factor model of spatial ability separating objects-based and orienting tests combining putatively separate categories of tests administered across the two batteries. ND = navigation based on directions, NL = navigation based on landmarks, MR = map reading, RM = route memory, PT = perspective taking, SC = scanning, CS = cross-sections, 2d = 2d drawing, PA = pattern assembly, EM =Elithorn Mazes, MR = Mechanical Reasoning, PF = paper folding, 3d = 3d drawing, Rot = mental rotation, PT = perspective taking, Maz = mazes; Model fit indices are reported in Table S5.

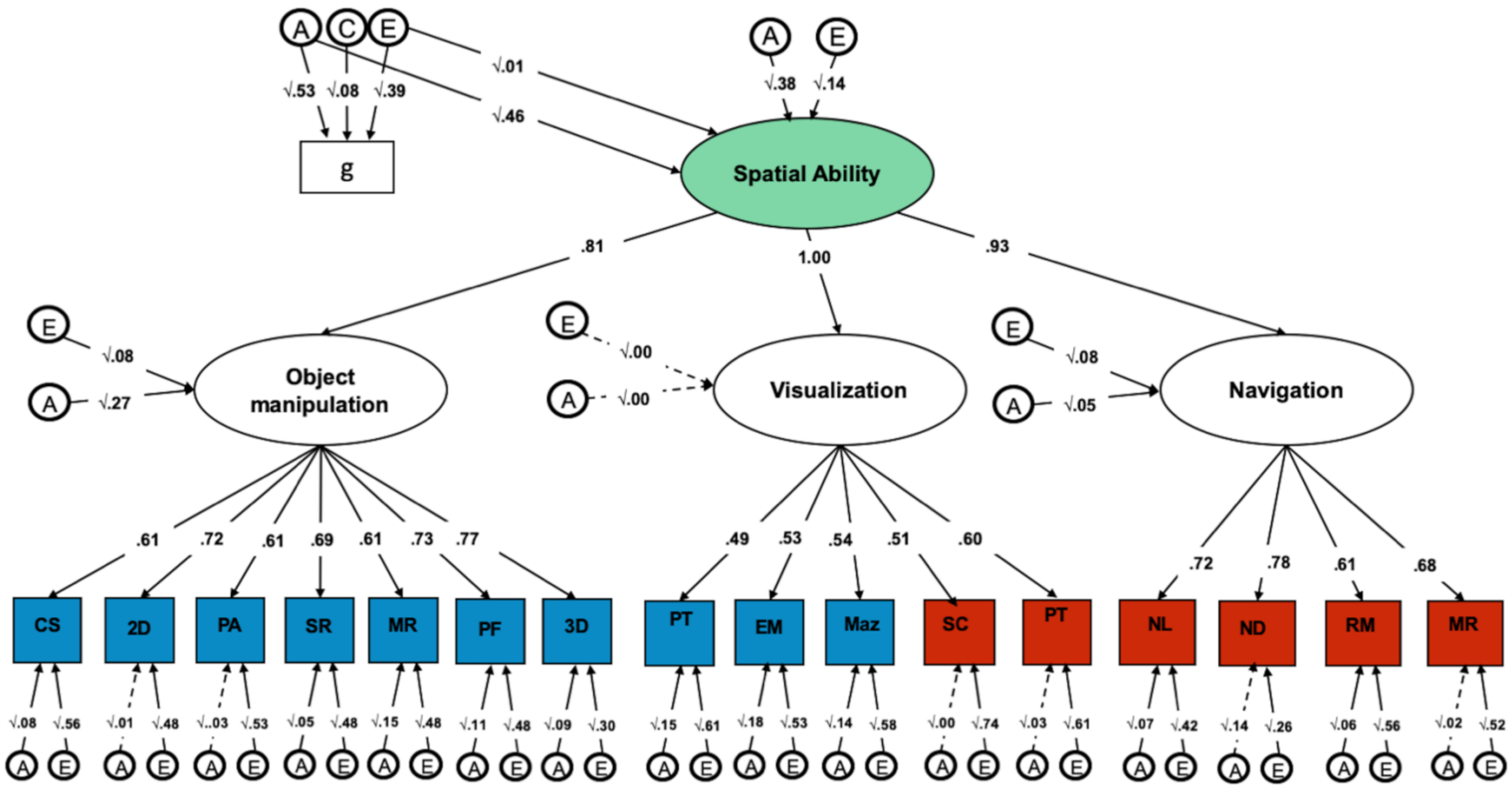


**Figure S3.** Hierarchical common pathway model exploring the genetic and environmental association between *g* and the common spatial ability factor. ND = navigation based on directions, NL = navigation based on landmarks, MR = map reading, RM = route memory, PT = perspective taking, SC = scanning, CS = cross-sections, 2d = 2d drawing, PA = pattern assembly, EM =Elithorn Mazes, MR = Mechanical Reasoning, PF = paper folding, 3d = 3d drawing, Rot = mental rotation, PT = perspective taking, Maz = mazes, *g* = general cognitive ability; Model fit indices are reported in Table S5.

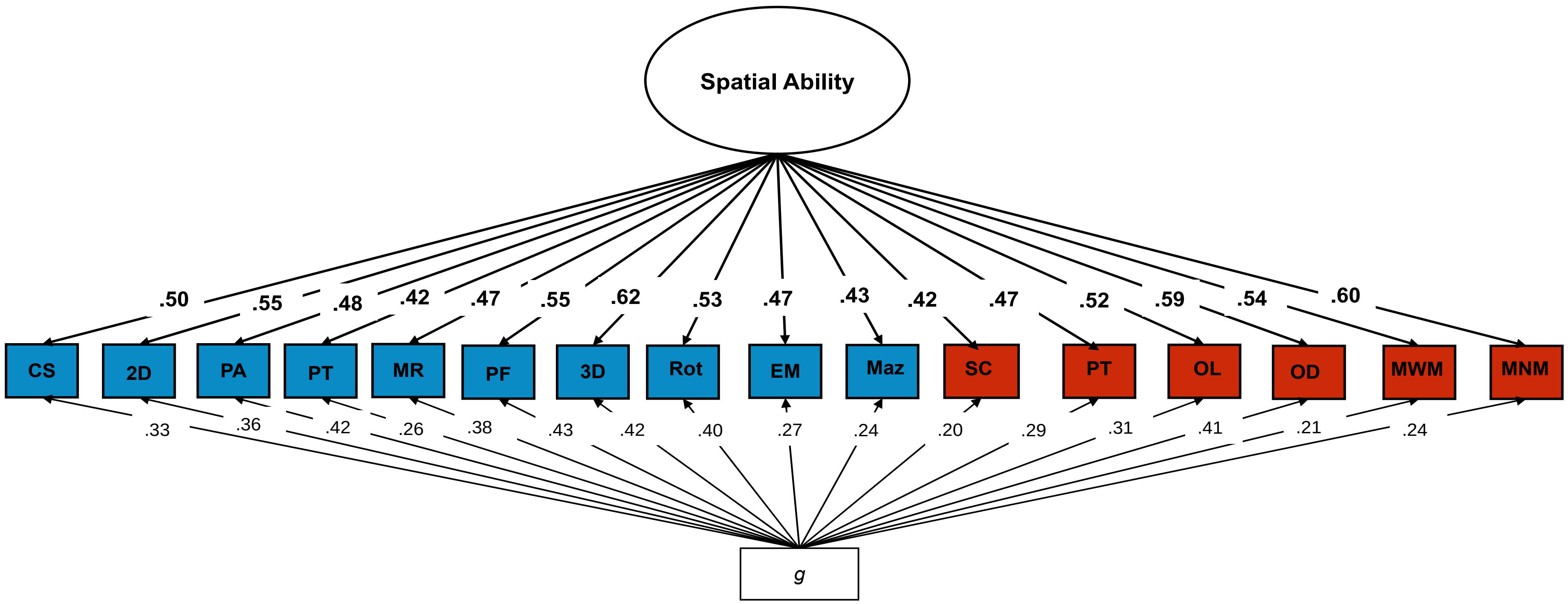


**Figure S4.** One-factor model of spatial ability including the 16 spatial tests accounting for *g* at the level of the indicator (each test); see Table S5 for model fit indices. ND = navigation based on directions, NL = navigation based on landmarks, MR = map reading, RM = route memory, PT = perspective taking, SC = scanning, CS = cross-sections, 2d = 2d drawing, PA = pattern assembly, EM = Elithorn Mazes, MR = Mechanical Reasoning, PF = paper folding, 3d = 3d drawing, Rot = mental rotation, PT = perspective taking, Maz = mazes, *g* = general cognitive ability.

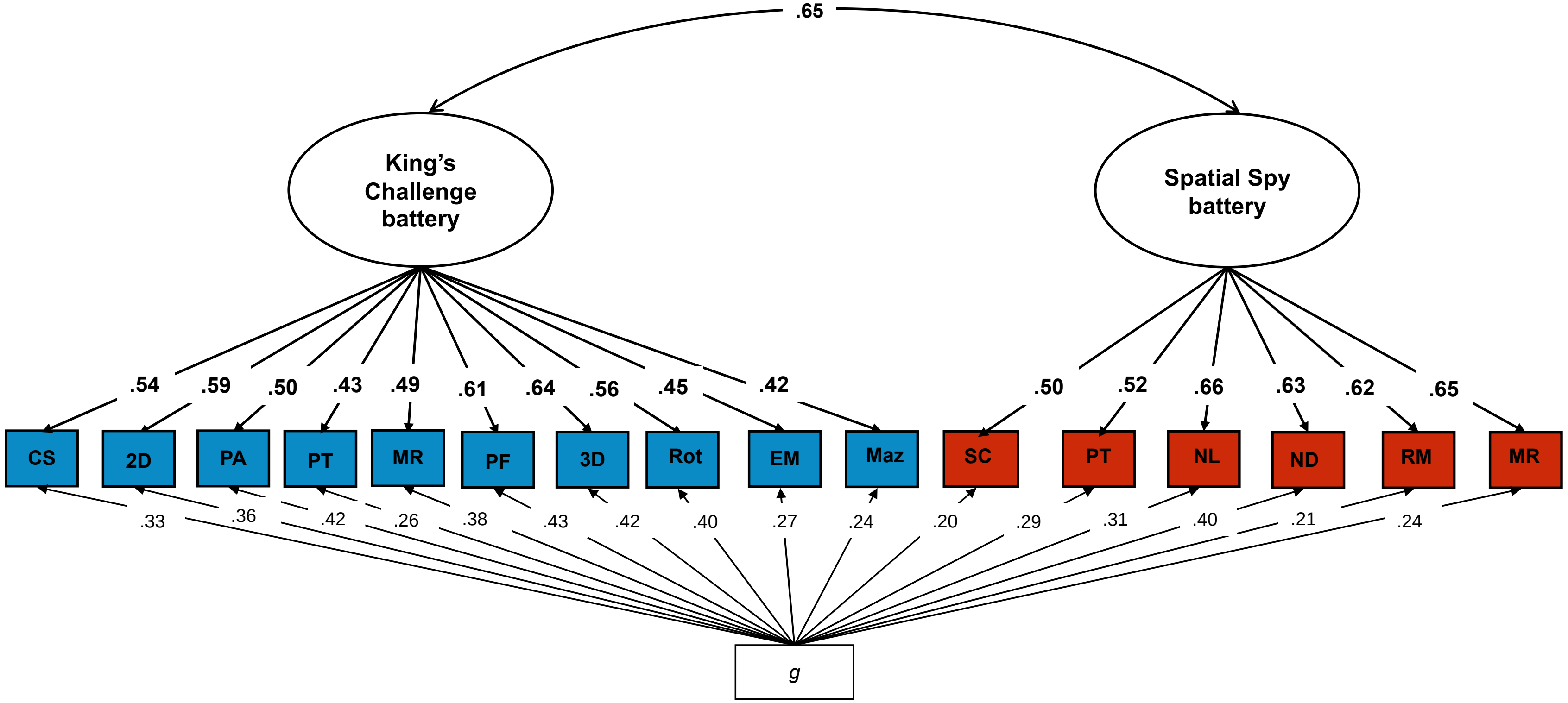


**Figure S5.** Two-factor model of spatial ability separating the two batteries accounting for *g* at the level of the indicators (each test). Model fit indices are reported in Table S5. ND = navigation based on directions, NL = navigation based on landmarks, MR = map reading, RM = route memory, PT = perspective taking, SC = scanning, CS = cross-section, 2d = 2d drawing, PA = pattern assembly, EM =Elithorn Mazes, MR = Mechanical Reasoning, PF = paper folding, 3d = 3d drawing, Rot = mental rotation, PT = perspective taking, Maz = mazes, *g* = general cognitive ability.

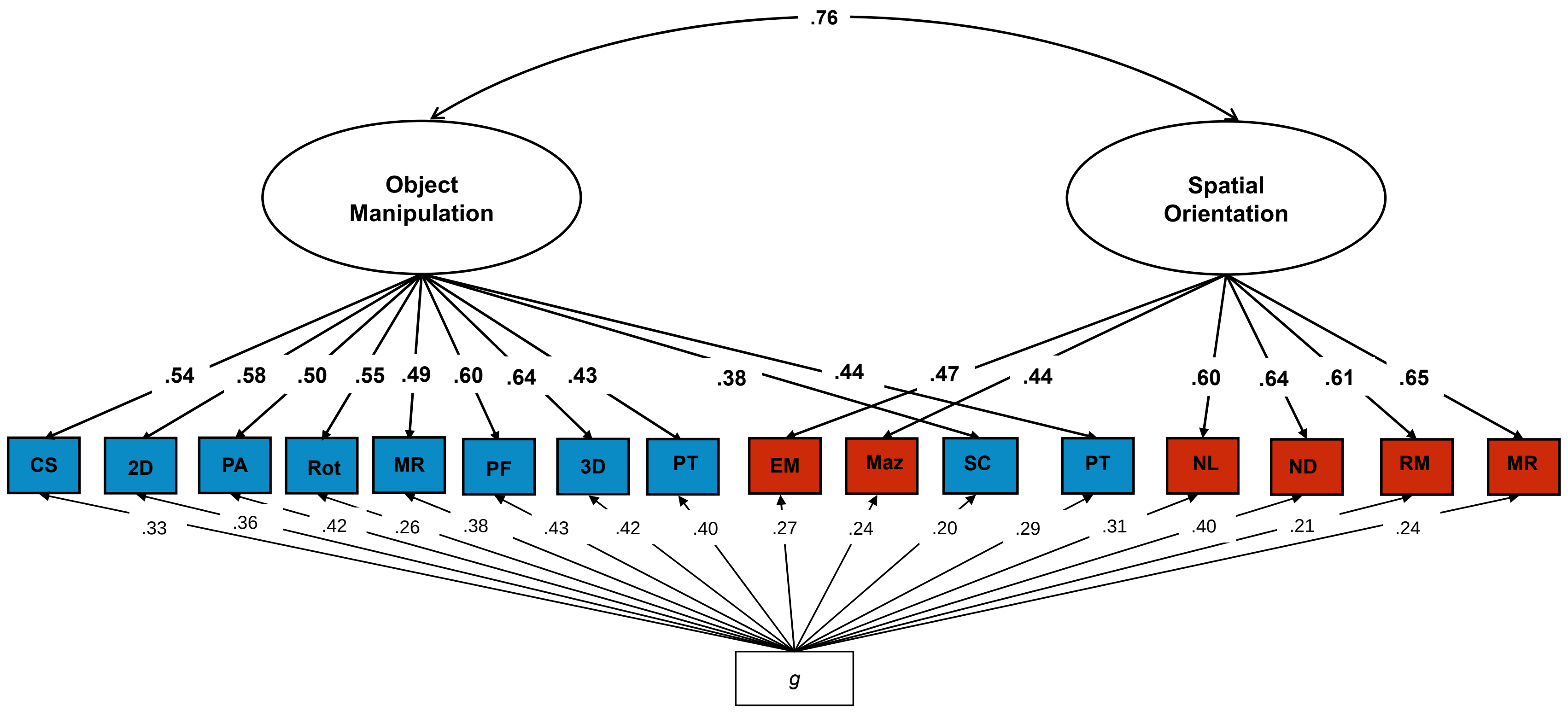


**Figure S6.** Two-factor model of spatial ability separating objects-based and orienting tests combining putatively different aspects of spatial skills across the two batteries accounting for *g* at the level of the indicators (each test). Model fit indices are reported in Table S5. ND = navigation based on directions, NL = navigation based on landmarks, MR = map reading, RM = route memory, PT = perspective taking, SC = scanning, CS = cross-sections, 2d = 2d drawing, PA = pattern assembly, EM =Elithorn Mazes, MR = Mechanical Reasoning, PF = paper folding, 3d = 3d drawing, Rot = mental rotation, PT = perspective taking, Maz = mazes, *g* = general cognitive ability.

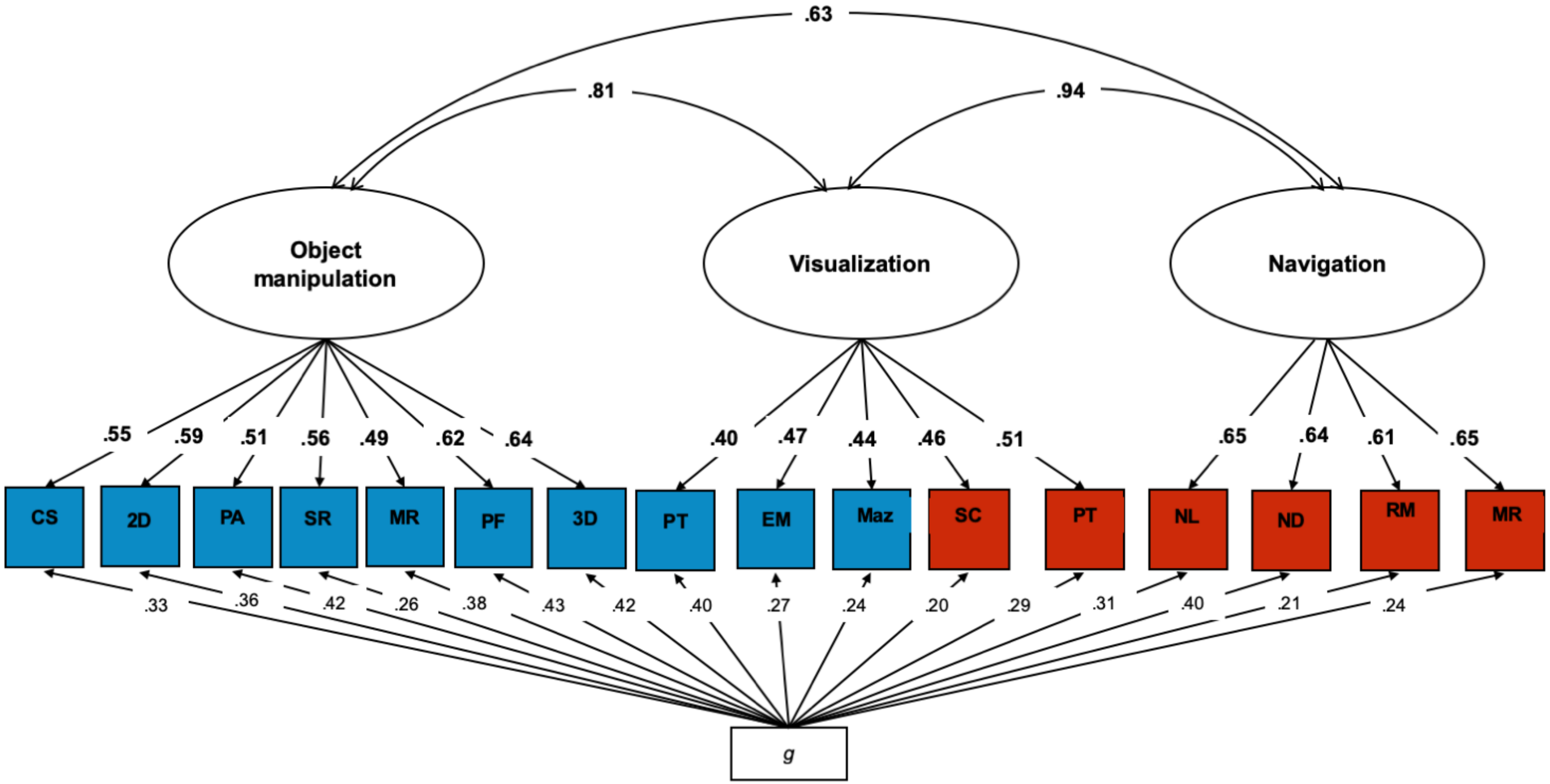


**Figure S7.** Three-factor model of spatial ability separating objects-based, navigation and visualization tests across the two batteries accounting for *g*. Model fit indices are reported in Table S5. ND = navigation based on directions, NL = navigation based on landmarks, MR = map reading, RM = route memory, PT = perspective taking, SC = scanning, CS = cross-sections, 2d = 2d drawing, PA = pattern assembly, EM =Elithorn Mazes, MR = Mechanical Reasoning, PF = paper folding, 3d = 3d drawing, Rot = mental rotation, PT = perspective taking, Maz = mazes, *g* = general cognitive ability.

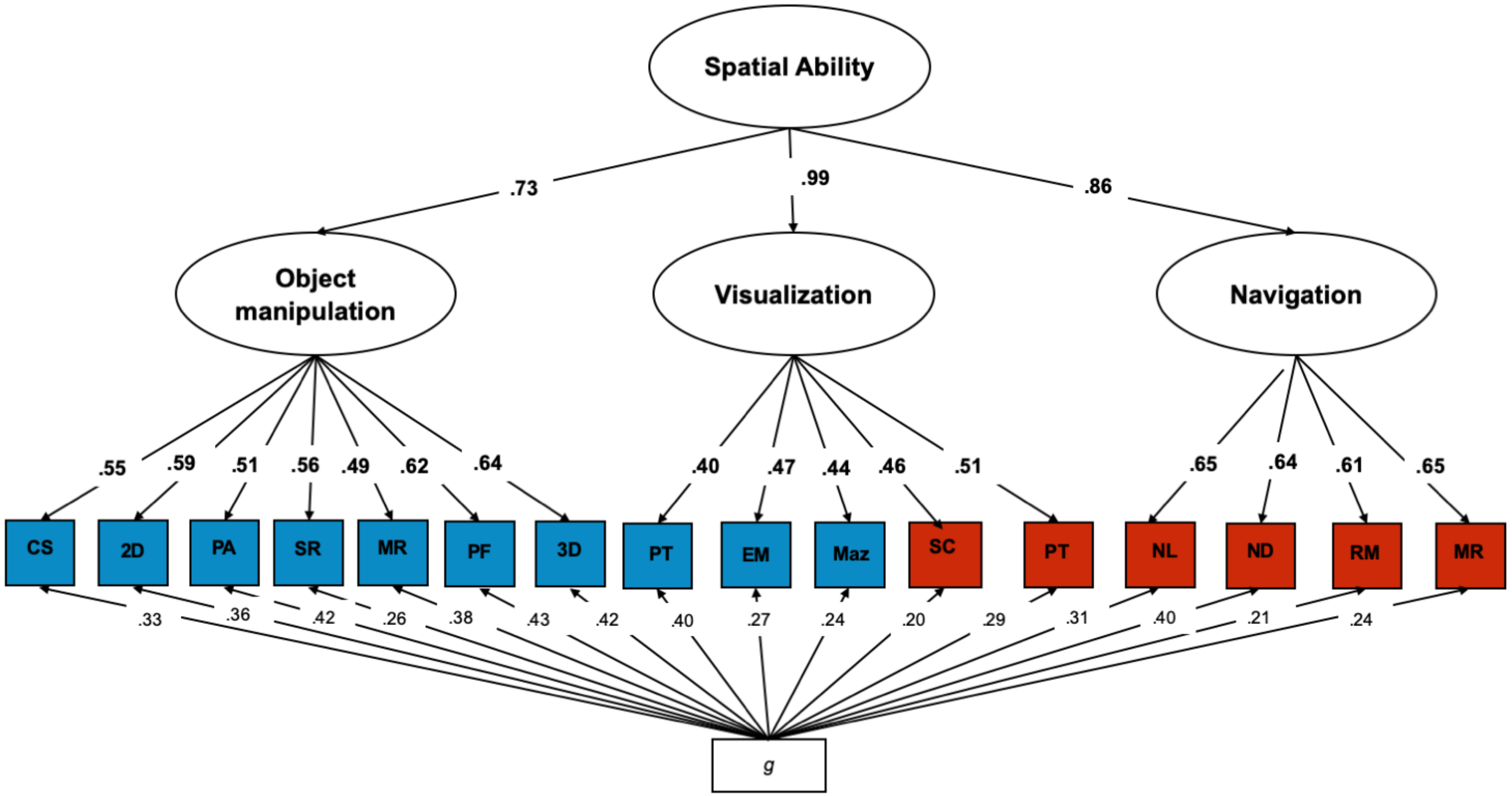


**Figure S8.** Hierarchical model including three first-order spatial factors (Navigation, Object-based and Visualization) and a second-order common factor of Spatial ability, accounting for *g* at the level of the indicators (each test). Model fit indices are reported in Table S5. ND = navigation based on directions, NL = navigation based on landmarks, MR = map reading, RM = route memory, PT = perspective taking, SC = scanning, CS = cross-sections, 2d = 2d drawing, PA = pattern assembly, EM =Elithorn Mazes, MR = Mechanical Reasoning, PF = paper folding, 3d = 3d drawing, Rot = mental rotation, PT = perspective taking, Maz = mazes, *g* = general cognitive ability.

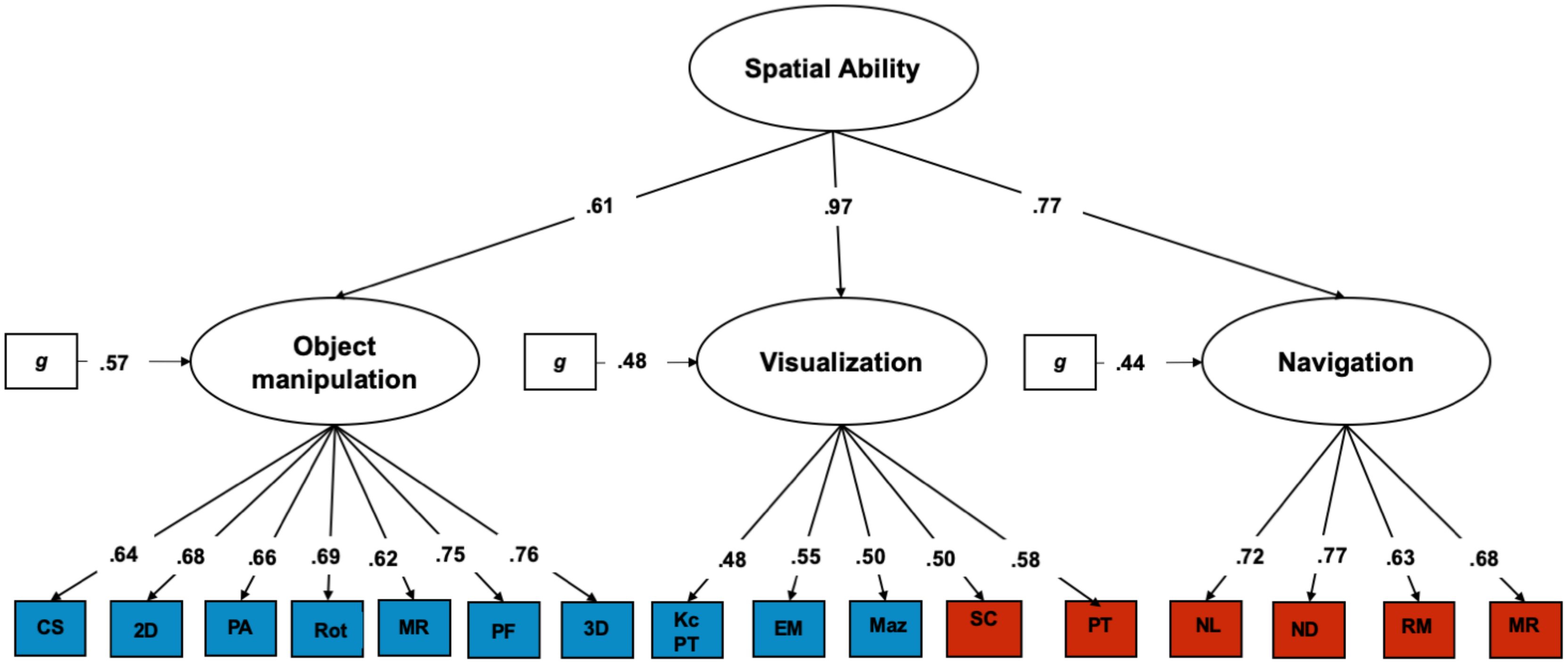


**Figure S9.** Hierarchical model including three first-order spatial factors (Navigation, Object-based and Visualization) and a second-order common factor of Spatial ability, accounting for *g* at the level of the first-order factors. Model fit indices are reported in Table S5. ND = navigation based on directions, NL = navigation based on landmarks, MR = map reading, RM = route memory, PT = perspective taking, SC = scanning, CS = cross-sections, 2d = 2d drawing, PA = pattern assembly, EM =Elithorn Mazes, MR = Mechanical Reasoning, PF = paper folding, 3d = 3d drawing, Rot = mental rotation, PT = perspective taking, Maz = mazes, *g* = general cognitive ability.

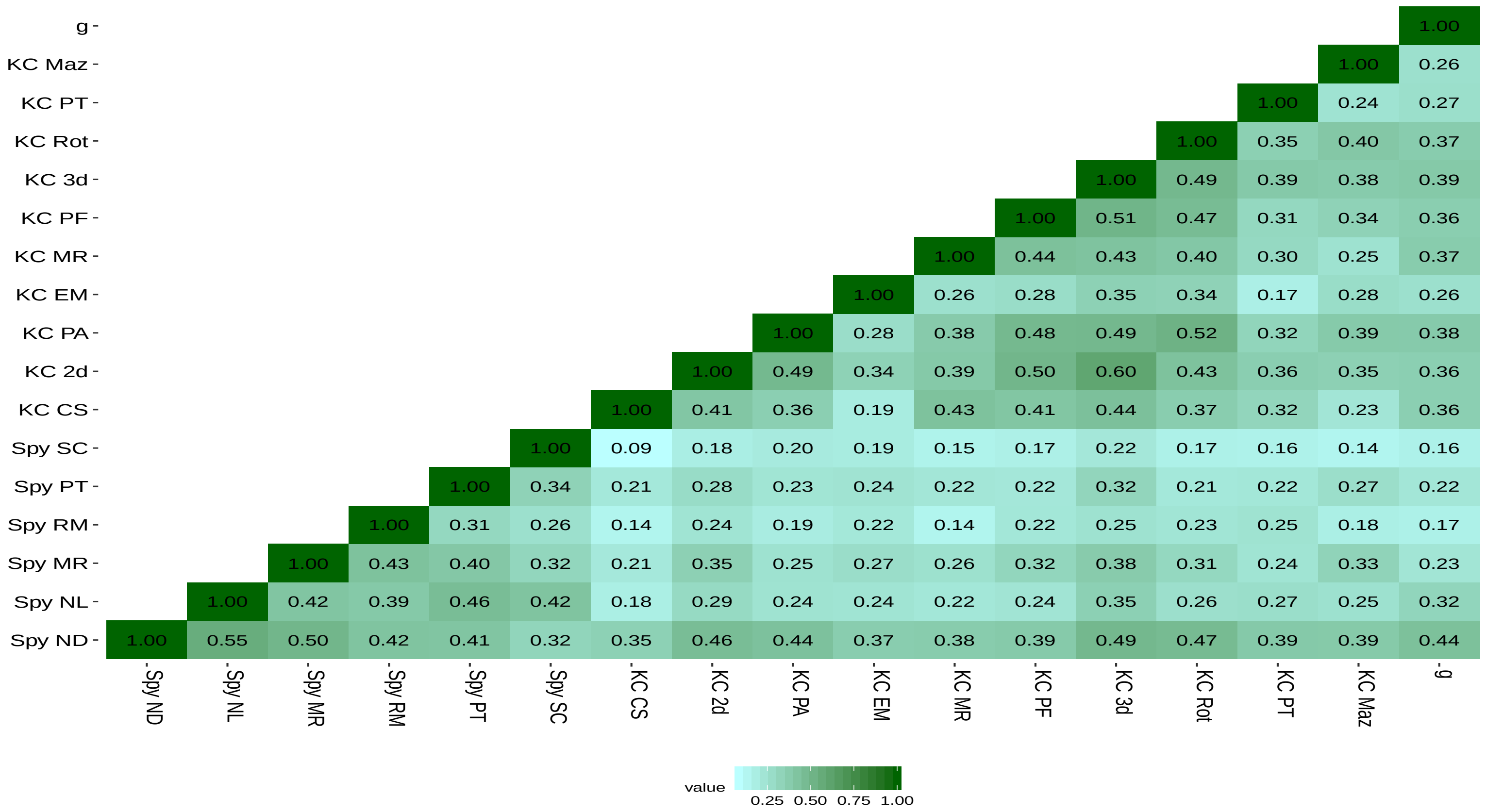


**Figure S10.** Correlations between all spatial tests and general cognitive ability using data from the other half of the phenotypic sample. Spy = Spatial Spy battery (large-scale), KC = King's Challenge battery (small-scale), ND = navigation based on directions, NL = navigation based on landmarks, MR = map reading, RM = route memory, PT = perspective taking, SC = scanning, CS = cross-sections, 2d = 2d drawing, PA = pattern assembly, EM =Elithorn Mazes, MR = Mechanical Reasoning, PF = paper folding, 3d = 3d drawing, Rot = mental rotation, PT = perspective taking, Maz = mazes, *g* = general cognitive ability. All correlations were significant at the $p < .001$ level; variables were residualized for age and sex and standardized prior to analysis.

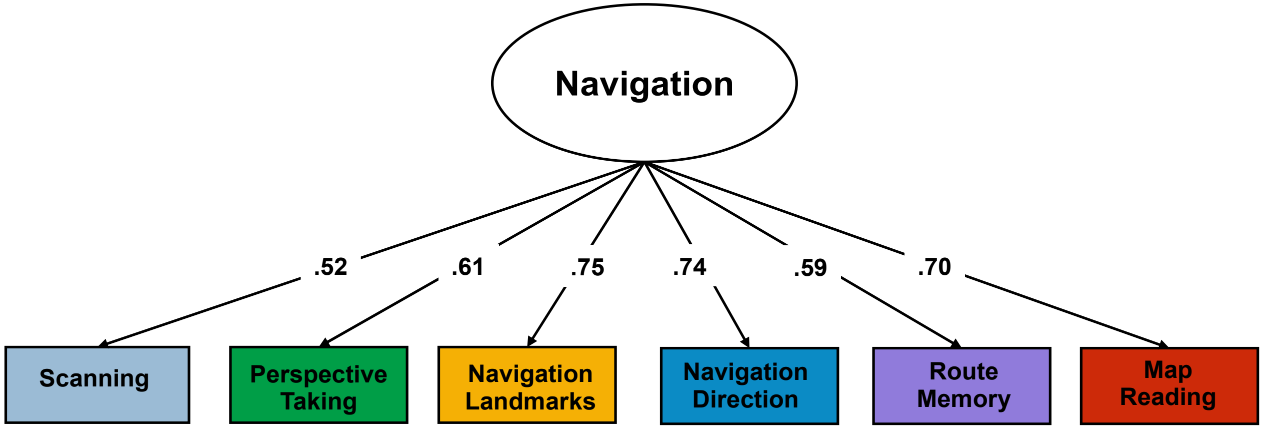


**Figure S11.** Factor structure of navigation abilities conducted examining the other half of the sample; CFI = 0.968, TLI = 0.947, RMSEA = 0.073, SRMR = 0.027.

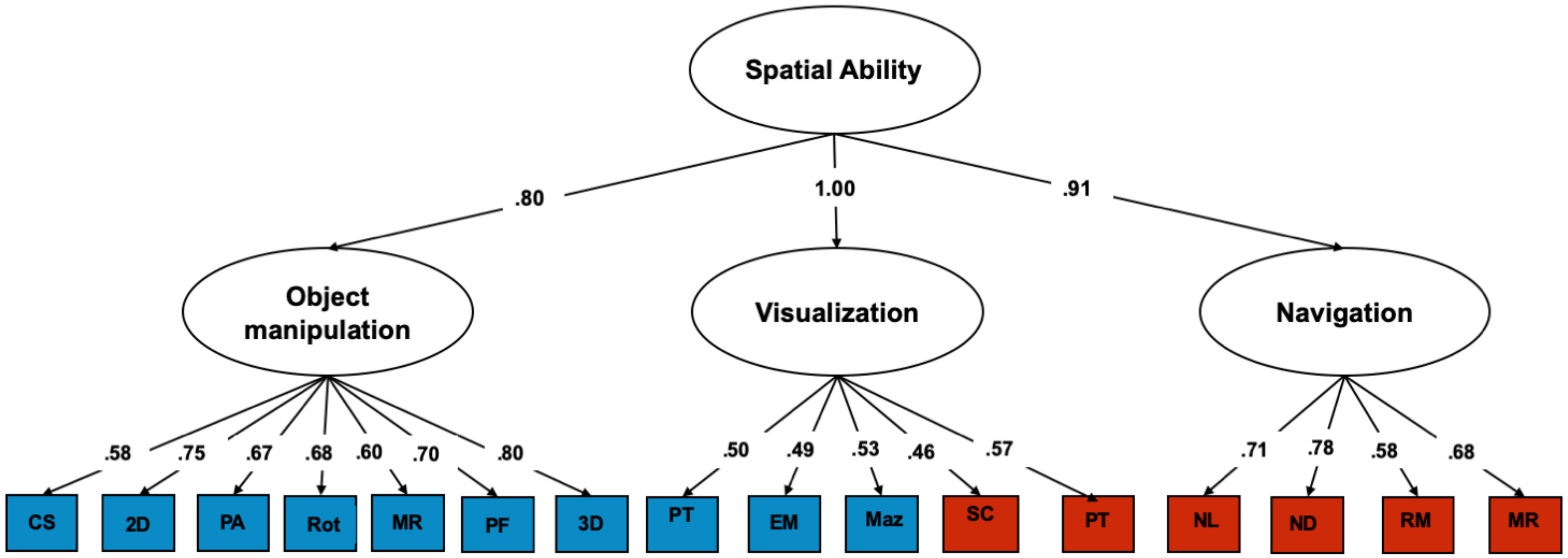


**Figure S12.** Hierarchical model of spatial abilities conducted in the other half of the sample; CFI = 0.939, TLI = 0.928, RMSEA = 0.046, SRMR = 0.050.
